# Accurate rare variant phasing of whole-genome and whole-exome sequencing data in the UK Biobank

**DOI:** 10.1101/2022.10.19.512867

**Authors:** Robin J. Hofmeister, Diogo M. Ribeiro, Simone Rubinacci, Olivier Delaneau

**Affiliations:** Department of Computational Biology, University of Lausanne, Lausanne, Switzerland

## Abstract

The UK Biobank performed whole-genome sequencing (WGS) and whole-exome sequencing (WES) across hundreds of thousands of individuals, allowing researchers to study the effects of both common and rare variants. Haplotype phasing distinguishes the two inherited copies of each chromosome into haplotypes and unlocks novel analyses at the haplotype level. In this work, we describe a new phasing method, SHAPEIT5, that accurately and rapidly phases large sequencing datasets and illustrates its key features on the UK Biobank WGS and WES data. First, we show that it phases rare variants with high accuracy. For instance, variants found in 1 sample out of 100,000 in the WES data are phased with accuracy above 95%. Second, we show that it can phase singletons, although with moderate accuracy, thereby making their inclusion in downstream analyses possible. Third, we show that the use of UK Biobank as a reference panel increases the accuracy of genotype imputation, an increase that is more pronounced when phased with SHAPEIT5 compared to other methods. Finally, we screen the phased WES data for loss-of-function (LoF) compound heterozygous (CH) events and identify 549 genes in which both gene copies are found knocked out. This list of genes complements current knowledge of gene essentiality in the human genome. We provide SHAPEIT5 in an open-source format, providing researchers with the means to leverage haplotype information in genetic studies.

## Introduction

Modern genetic association studies are based on whole-genome or whole-exome sequencing (WGS/WES) for hundreds of thousands of samples, data usually collected as part of nationwide biobanking initiatives^1,2^. Compared to previous studies that were mostly based on SNP arrays, WGS and WES data can identify rare variants, allowing a systematic characterization of their contribution to trait heritability^3^, their functional relevance^4^ and their effects on various traits and diseases^5,6^. In this context, haplotype phasing of rare variants, which involves distinguishing the two parentally inherited copies of each chromosome into haplotypes, adds an additional layer of biologically relevant information and unlock novel analyses. For instance, phasing is crucial to identify compound heterozygous (CH) events, which occur when both copies of a gene contain heterozygous mutations. In the case of Mendelian disorders, CH is known to be one of the most common inheritance models for rare recessive diseases in non-consanguineous individuals^7,8^. However, the contribution of CH to complex traits and diseases remains to be characterized in large cohorts. This requires phase information to be retrieved with high confidence and to be considered when rare variants are analyzed, such as in gene-based burden test analysis^9^. The most reliable approach to phase rare variants in large cohorts of individuals is statistical phasing, which leverages information across individuals to make estimation^10^. This technique is well established for common variants typed on SNP arrays, where phase information is used, for instance, to perform genotype imputation^11^, admixture analysis^12^ and genealogy estimation^13^. Phasing methods have been heavily optimized to scale efficiently to the thousands of samples in modern SNP array datasets, but not to the dozens of millions of rare variant sites present in WGS/WES datasets. As an example, the WGS data for 150,119 UK Biobank samples comprises three orders of magnitude more variants than the Axiom array data, ∼96% of them having a minor allele frequency (MAF) below 0.1%. Recently, a solution to this problem has been proposed^14,15^, in which common and rare variants are phased separately: in a first step a standard phasing method is used to obtain haplotypes at common variants, and in a second step rare heterozygous sites are phased onto the resulting haplotypes using genotype imputation technique. This type of strategy, based on building first a scaffold of haplotypes, has been used in other contexts, such as in genotype imputation^16^, integration of family data^17^ and integration of external phasing information^18^.

In this work, we describe a new phasing method, SHAPEIT5, designed to accurately phase rare variants in large WGS/WES datasets, including singletons with moderate accuracy, while attributing phasing confidence scores. We applied our method to estimate accurate haplotypes for 150,119 UK Biobank samples with WGS data and 452,644 UK Biobank samples with WES data. We then demonstrate the benefit of using these two haplotype collections as reference panels for SNP array imputation and finally show that the phase inferred at rare variants in the WES dataset can be screened to reliably identify compound heterozygous loss-of-function (LoF) mutations, likely leading to complete gene knockouts.

## Results

### Overview of the SHAPEIT5 phasing method

SHAPEIT5 performs haplotype phasing of WGS or WES data using three different phasing models, each focussing on a specific type of variants: (i) common variants are phased using the SHAPEIT4 model^18^, (ii) then rare variants are phased onto the resulting haplotypes using an imputation model and (iii) finally, singletons are phased using a simple coalescent-inspired model. See **Figure 1** for an illustration of the overall phasing scheme. Common variants are defined here as having a minor allele frequency (MAF) above 0.1% and are phased using an optimized version of the SHAPEIT4 algorithm^18^, already known to perform well on large sample size (**Figure 1a**).

**Figure 1:**
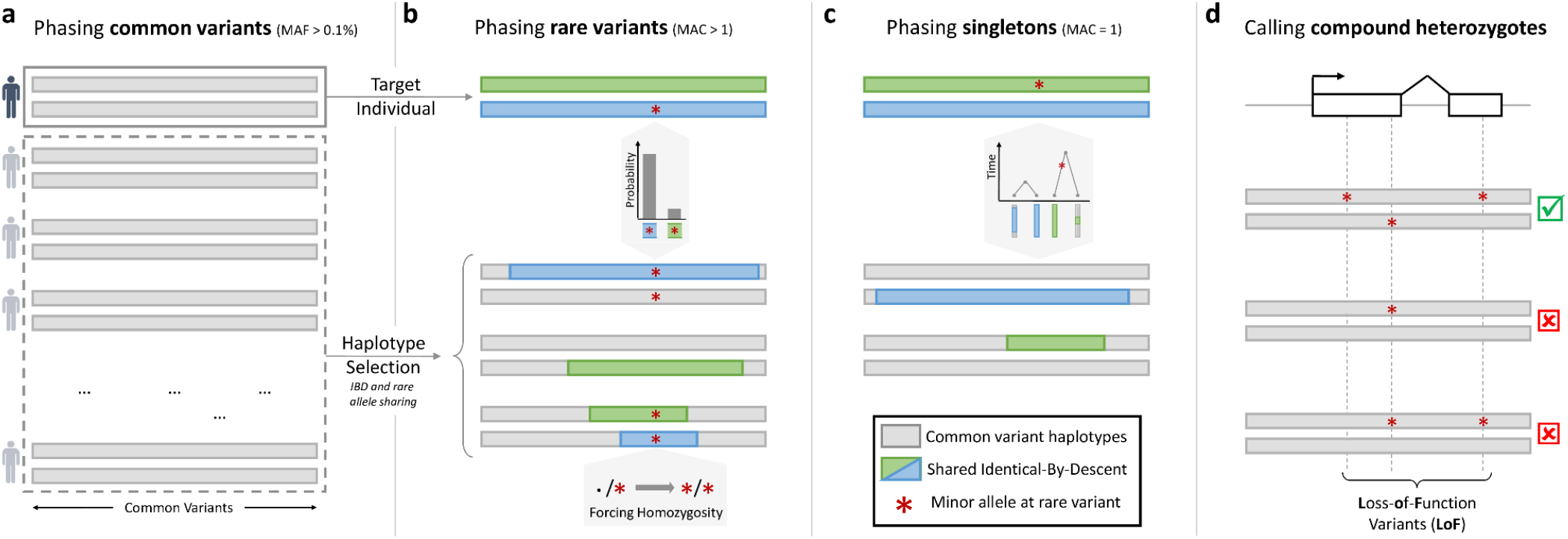
Rationale of SHAPEIT5. From left to right. (**a**) All samples are phased at common variants (MAF ≥ 0.1%). (**b**) Phasing of a given rare variant onto the haplotypes at common variants. Conditioning haplotypes used in the estimation share long matches with the target (in green and blue) and are not monomorphic at the rare variant. (**c**) Singleton phasing by assigning the new allele on the target haplotype with the shortest match. (**d**) Compound heterozygous event mapping based on the rare variant phasing (a-c).

Once the first phasing stage is completed at common variants, the resulting haplotypes are used in a second stage as a scaffold onto which rare variants (MAF < 0.1%) are phased one after another. To cope with the large numbers of rare variants, SHAPEIT5 uses a sparse data representation: only genotypes carrying at least one copy of the minor allele are stored in memory and considered for computation, thereby discarding all genotypes being homozygous for the major allele^15,19,20^. SHAPEIT5 phases each rare heterozygous genotype conditioning on a small number of informative haplotypes in the dataset (**Figure 1b**). For a specific rare variant, these conditioning haplotypes are chosen so that (1) they belong to samples being locally identical-by-descent (IBD) with the target sample and (2) they are not monomorphic at the rare variant (i.e. at least a few carry a copy of the minor allele). To comply with the first requirement, SHAPEIT5 uses a Positional Burrows-Wheeler Transform (PBWT) data structure^21^ built on all the scaffold haplotypes in the dataset. This allows rapid identification of long shared segments between haplotypes and therefore of IBD between individuals. In order to ensure representation of the minor allele in the conditioning set (second requirement), the method performs a second PBWT selection pass restricted to the subset of samples carrying a copy of the minor allele. This second pass is performed efficiently by leveraging the sparse representation of the genotypes. We then determine the alleles carried by the conditioning haplotypes at the rare variant of interest, which is straightforward when homozygous. However, when a conditioning sample is heterozygous, the allele carried by each of its two haplotypes is unknown. In this case, our model assumes that both haplotypes carry the minor allele^15^. Once the conditioning set of haplotypes is assembled, SHAPEIT5 uses the Li and Stephens model^22^ to get the most likely phase configuration of the rare allele by imputation (i.e. either on its first or second target haplotype; **Supplementary Fig. 1**). The strength of our model resides in the guarantee that each rare heterozygous genotype is phased from a conditioning set that contains long haplotype matches and that carries copies of the two possible alleles.

However, this approach is not possible for singletons (minor allele count of 1) as no other sample carries the minor allele. In this case, SHAPEIT5 uses another phasing model that (i) assumes singletons to be recent mutation events and (ii) leverages IBD sharing patterns between haplotypes to make inference (**Figure 1c**). Specifically, our model identifies the longest possible match in the dataset for each target haplotype. By definition, these matches point to haplotypes sharing recent common ancestors with the target and their lengths indicate the number of generations separating them: the shorter the match, the older the common ancestor. Our model assumes that an older common ancestor means more time for a mutation to occur on that lineage and therefore assigns the minor alleles of singletons to the target haplotype with the shortest match.

### Phasing UK Biobank exomes and genomes

We used SHAPEIT5 to phase haplotypes for three different sequencing datasets from the UK Biobank. The first dataset comprises the whole-genome sequencing (WGS) data on chromosome 20 for 147,754 samples and ∼13.8 million SNPs and indels after quality control. The second dataset comprises the whole-exome sequencing (WES) data for 452,644 samples and ∼26 million variants. In these two datasets, we only included samples for which Axiom array data is available and excluded parental genomes for duos (parent-offspring pairs) and trios (parent-offspring triplets) to measure phasing accuracy in the offsprings. The last dataset comprises the WGS data for the full set of 150,119 samples and ∼603 million variants. Numbers of samples, trios, duos and variants after quality control are given in **Supplementary Table 1**. Phasing of the WES dataset was performed for each chromosome independently and phasing of the WGS was done in overlapping chunks of ∼4.5Mb on average to leverage parallelization on the Research Analysis Platform (RAP) of the UK Biobank. To compare the performance of our method against other methods, we used Beagle5.4^15^ to phase the full WES dataset and the WGS dataset on chromosome 20, run with default parameters.

### Phasing performance in the UK Biobank data

To assess phasing performance, we used the available white British trios (719 for WES, 31 for WGS). As the 31 trios with WGS data are insufficient to reliably estimate phasing accuracy, we also considered 432 white British duos for the WGS dataset. Using these, (i) we first derived a true set of haplotypes for the offsprings using basic inheritance logic, (ii) we then performed statistical phasing of the WES and WGS datasets after having excluded parental genomes and (iii) we compared the offspring haplotypes obtained by statistically inference to the true set obtained in (i). We assessed how close the two sets of haplotypes are by measuring the switch error rate (SER), which is the fraction of successive heterozygous genotypes phased differently between the two sets. When looking at overall SER using different validation sets (duos, trios), different sets of variants (all variants or common variants only) and different sample sizes, we only found minor differences between SHAPEIT5 and Beagle5.4^15^ on the WGS data (**Supplementary Fig. 2a-c**). However, when only considering Axiom array positions, lower SER is observed with SHAPEIT5 (**Supplementary Fig. 2d**). Interestingly, we did not find the same pattern when phasing the Axiom array data only (n=5,000 to n=480,000). In this case, the two methods exhibit similar accuracy regardless of sample size (**Supplementary Fig. 3**). On this data, we obtained low SER (<0.2%) on the highest sample sizes for both methods, to the point that it becomes difficult to distinguish switch errors from genotyping errors (**Supplementary Fig. 4**).

A key feature of the WES and WGS datasets is the large number of rare variants they contain (MAF < 0.1%). As the number of heterozygous genotypes is low at these variants, they have a small contribution towards global SER measurements. We therefore stratified the SER within bins of minor allele counts (MAC) to evaluate the method performance on rare variants. In practice, we assigned heterozygous genotypes to different MAC bins depending on the variant frequency and we computed in each MAC bin the fraction of them being correctly phased (relative to the previous heterozygous genotype). When doing so, we found that rare variants are phased by SHAPEIT5 with higher accuracy than Beagle5.4 in both the WGS and WES datasets (**Figures 2a-b**). For instance, extremely rare variant sites in the WGS data (MAC between 11 and 20) are phased by SHAPEIT5 with a switch error rate of 4.36%, which compares to 8.76% when phased by Beagle5.4. This represents a drop of 50.2% in SER compared to Beagle5.4. Similarly, in the WES callset, the same variant category is phased by SHAPEIT5 with a switch error rate of 2.93% compared to 5.18% when using Beagle5.4 (42.67% reduction). Overall, SHAPEIT5 phases rare variants in the WES and WGS with 20% to 50% fewer switch errors compared to Beagle5.4, depending on MAC. Importantly, this improvement in accuracy is also observed when validating haplotypes only using trios in the WGS dataset (**Supplementary Fig. 5**) and is sample-size dependent (**Supplementary Fig. 6**). Indeed, significant differences between the two methods are observed from WGS datasets comprising at least 50,000 samples and the differences increase as sample size increases.

**Figure 2:**
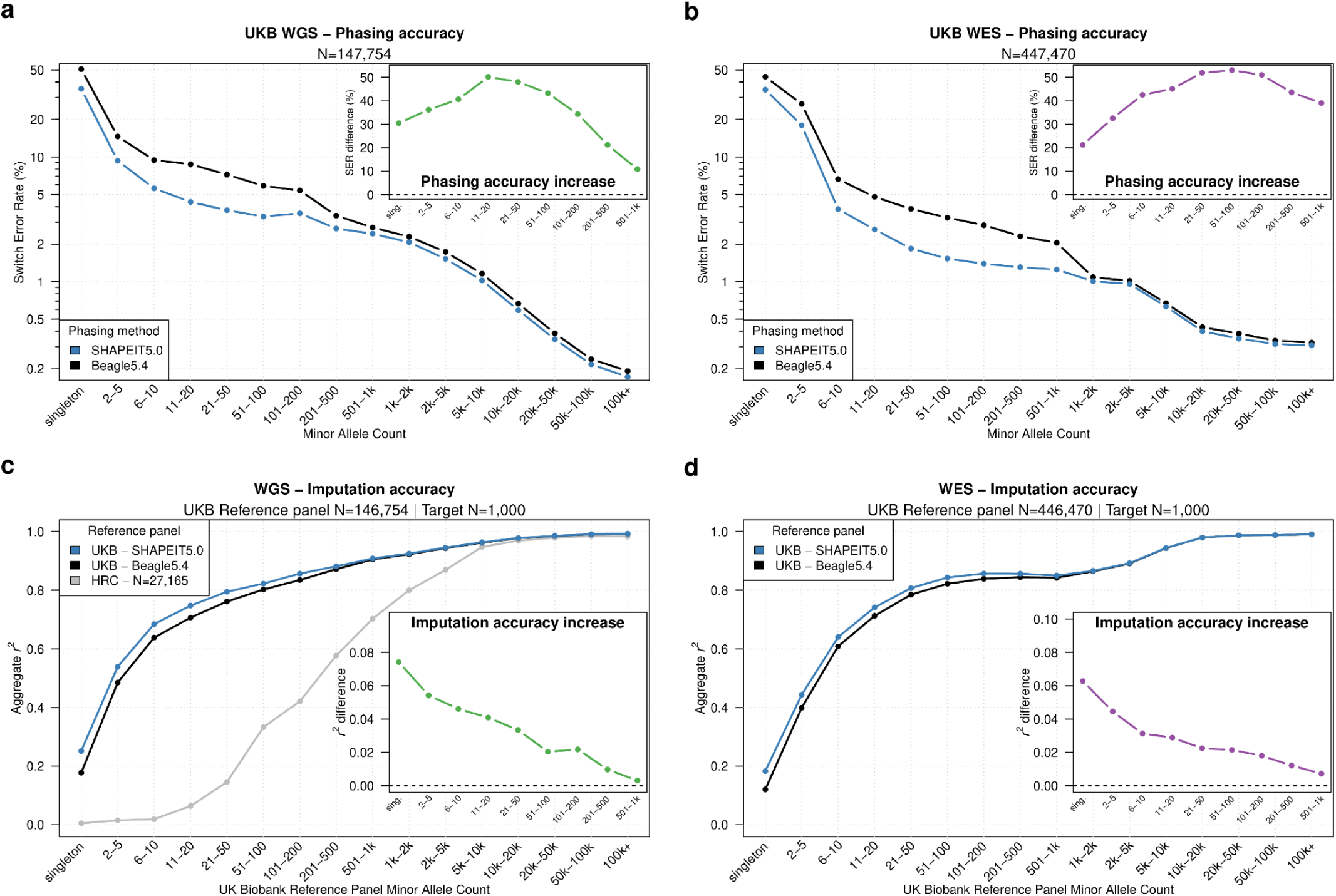
Phasing performance. (**a**,**b**) Switch error rate (SER, y-axis, log-scale) of SHAPEIT5 (blue) compared to Beagle5.4 (black) stratified by minor allele count (x-axis) for the UK Biobank whole genome sequencing (**a**) and whole exome sequencing (**b**). The zoomed-in views show the relative reduction of switch error rate using SHAPEIT5 compared to Beagle5.4 at rare variants. (**c**,**d**) Imputation accuracy (Aggregate r^2^, y-axis, log-scale) for 1,000 white British samples genotyped with the Axiom array when using reference panels phased with either SHAPEIT5 (blue) or Beagle5.4 (black) whole genome sequencing (**c**) or whole exome sequencing (**d**). In (**c**) the data was also imputed using the HRC reference panel (gray). The zoomed-in views show the increase of imputation accuracy at rare variants using the UK Biobank dataset phased with SHAPEIT5 compared to Beagle5.4 as a reference panel.

One novel feature of SHAPEIT5 is to provide phasing for singletons, which cannot be phased using previous population-based methods. In a large sequencing dataset, a singleton can be the product of multiple causes, including recent mutation, *de novo* mutation, somatic mutation or genotyping error. The model we propose aims at resolving the phase of recent mutations. We therefore estimated the fraction of singletons falling in this specific category by assessing duos and trios in the WGS data. Specifically, we measured the fraction of singletons in offspring that is not supported by the genotype data available for the parents. In duos, we found that 47.36% of the singletons are supported by the genotyped parent, whereas 52.64% are not (**Supplementary Fig. 7a**), suggesting that 5.26% of the singletons are not inherited from parents (assuming no inheritance bias). Consistently, we found that 4.52% of the singletons in trio offsprings are not inherited from the parents (none of the parents carry the minor allele; Mendel inconsistency; **Supplementary Fig. 7b**). Together, this shows that the vast majority of the singletons (∼95%) are inherited and can therefore be phased using both inheritance logic in trios and duos and our new model. In the WGS dataset, we obtained SER of 35.1% and 36.6% using duo and trio phasing as ground truth, respectively (**Supplementary Fig. 7c-d**). In the WES dataset, we obtained a SER of 35.2% using trio phasing as truth (**Figure 2b**). While relatively high, this is a significant deviation from the expected 50% from other methods (binomial test p-values < 2.2e-16, **Supplementary Fig. 7c-d**).

All computations have been performed on the Research Analysis Platform (RAP) of the UK Biobank. The RAP offers a choice of two priority levels for computations, “spot” (lower cost) and “on-demand” (higher cost). Assuming that all computing is performed on-demand, Beagle5.4 and SHAPEIT5 require 57.8 and 74.9 British pounds of computing costs (as of October 2022) to phase chromosome 20 with WGS data (147,754), which corresponds to approximately 2,842 and 3,746 British pounds for the entire genome (**Supplementary Table 2**). However, these are conservative estimates, as SHAPEIT5 allows phasing of the data in chromosomal chunks (in parallel), therefore greatly reducing the need of using “on-demand” priority compared to Beagle5.4 (**Supplementary Table 2**).

### SHAPEIT5 phasing improves the accuracy of genotype imputation

Several downstream analyses in disease and population genetics require haplotype-level data. One example is genotype imputation^23^, which uses large cohorts of WGS data as a reference panel to predict missing genotypes of SNP array data. As the accuracy of genotype imputation depends on the reference panel, we aim to quantify the impact of phasing errors on genotype imputation. This has two major advantages. First, it provides an alternative validation of the haplotypes that does not rely on SER and is easy to partition by minor allele frequency. Second, it assesses the quality of the phasing across all samples globally, and not only on a small subset for which parental genomes are available. Therefore, we imputed a subset of 1,000 British samples from the UK Biobank dataset from which we have SNP array data available, together with WGS and WES that we can use as validation.

First, we show that genotype imputation using the UKB WGS reference panel massively outperforms the previous generation of reference panels, such as the HRC^24^ (**Figure 2c)**, in line with previous findings that showed that large WGS panels can enhance imputation of genotyped samples^2^. For both UKB WGS and WES, we find that the reference panels phased with SHAPEIT5 outperform the reference panels phased with Beagle5.4 at rare variants (MAC < 500; **Figure 2c-d**), in agreement with the switch error rate estimates reported in **Figure 2a-b**. As an example, imputation using the WGS or WES reference panel phased with SHAPEIT5 provides an increase of squared Pearson coefficient of ∼0.05 for variants with a MAC between 2 to 5 compared to imputation performed on reference panels phased with Beagle5.4. In an association study, this corresponds to an increase of 5% in effective sample size when testing these variants for association, only due to better phasing of the reference panel^25^. Interestingly, even variants with a minor allele count of 1 in the reference panel are better imputed using the SHAPEIT5 reference panel, thereby confirming the validity of our phasing at singletons. When looking at the non-reference discordance (NRD, see Methods) of imputed genotypes partitioned by minor allele frequency, imputation from the SHAPEIT5 reference panels can reduce NRD up to 5.9% for WGS data and 5.3% for WES data compared to Beagle5.4 (**Supplementary Fig. 8**).

An additional feature of SHAPEIT5 is the introduction of a metric of phasing confidence at rare heterozygous sites (MAF<0.1%). This metric allows controlling for phasing errors and utilizing the certainty of the probabilistic phasing inference in downstream analyses, as in genotype imputation. The phasing confidence is designed to lie in the range of 0.5 to 1, where a value of 1 indicates that there is no uncertainty in the phase of the heterozygous genotype whereas a value of 0.5 means that the possible phases are equally likely. All singletons are attributed a phasing confidence of 0.5 as they are phased with a different model from which the confidence cannot be estimated. We assessed the phasing accuracy at different phasing confidence score thresholds (**Supplementary Fig. 9**). We show that filtering variants with a phasing confidence score threshold of 0.9 controls the switch error rate to a maximum of ∼2% for WGS data and ∼1% for WES data (with >80% and >50% variants retained), allowing downstream applications to confidently use rare and ultra rare heterozygous genotypes in the analysis.

### Identification of loss-of-function compound heterozygous mutations

Compound heterozygous (CH) events occur in an individual when both copies of a gene contain at least one heterozygous variant. Compound heterozygosity is often studied in the context of loss-of-function (LoF) variants, which are expected to have a highly deleterious effect on a gene – a rare event equivalent to a homozygous gene knockout. Indeed, CH events have been linked to multiple diseases including cancer, birth defects and Alzheimer’s disease^8,26–29^. The accurate haplotype phasing across the UK biobank performed in this study, including extremely rare variants, allows the identification of individuals and genes with CH events. To focus on potentially high phenotypic effects, we gathered 383,637 high-confidence loss-of-function (LoF) variants (stop-gain, frameshift or essential splice variants) across 374,826 white British individuals and 17,689 protein-coding genes phased with SHAPEIT5 (see Methods). We found that a gene has on average 22.3 LoF variants across the cohort and an individual has on average 7.7 LoF variants (**Supplementary Fig. 10**). To determine CH events, we identify individuals with LoF mutations in both copies of a gene. A total of 3,018 (17%) out of 17,689 genes tested had at least one individual with at least two LoF variants. From those 3,018 genes, we found 574 (19%) genes with one or more individuals with CH LoF variants (**Figure 3a**), for a total of 829 gene-individual CH events (816 distinct individuals, **Supplementary Fig. 11, Supplementary Data 1**). When considering only high-confidence haplotype calls (phasing confidence score > 0.99 and excluding singletons), we still identify 77% (441) genes and 74% (614) of the CH events identified in the full dataset, indicating that the majority of the CH events identified rely on high confidence haplotype calls (**Figure 3a, Supplementary Fig. 11**). We found that the 574 CH genes are highly depleted in several lists of known essential genes, compared to the 3,018 genes with two or more LoF variants (OR 0.1-0.49, p-value < 2.4e^-3^, **Figure 3b**). Conversely, CH genes are enriched in lists of non-essential and homozygous LoF tolerant genes (OR 1.28-3.10, **Figure 3c**). As the UK Biobank is largely composed of healthy individuals, a depletion of CH events in essential genes is expected.

**Figure 3:**
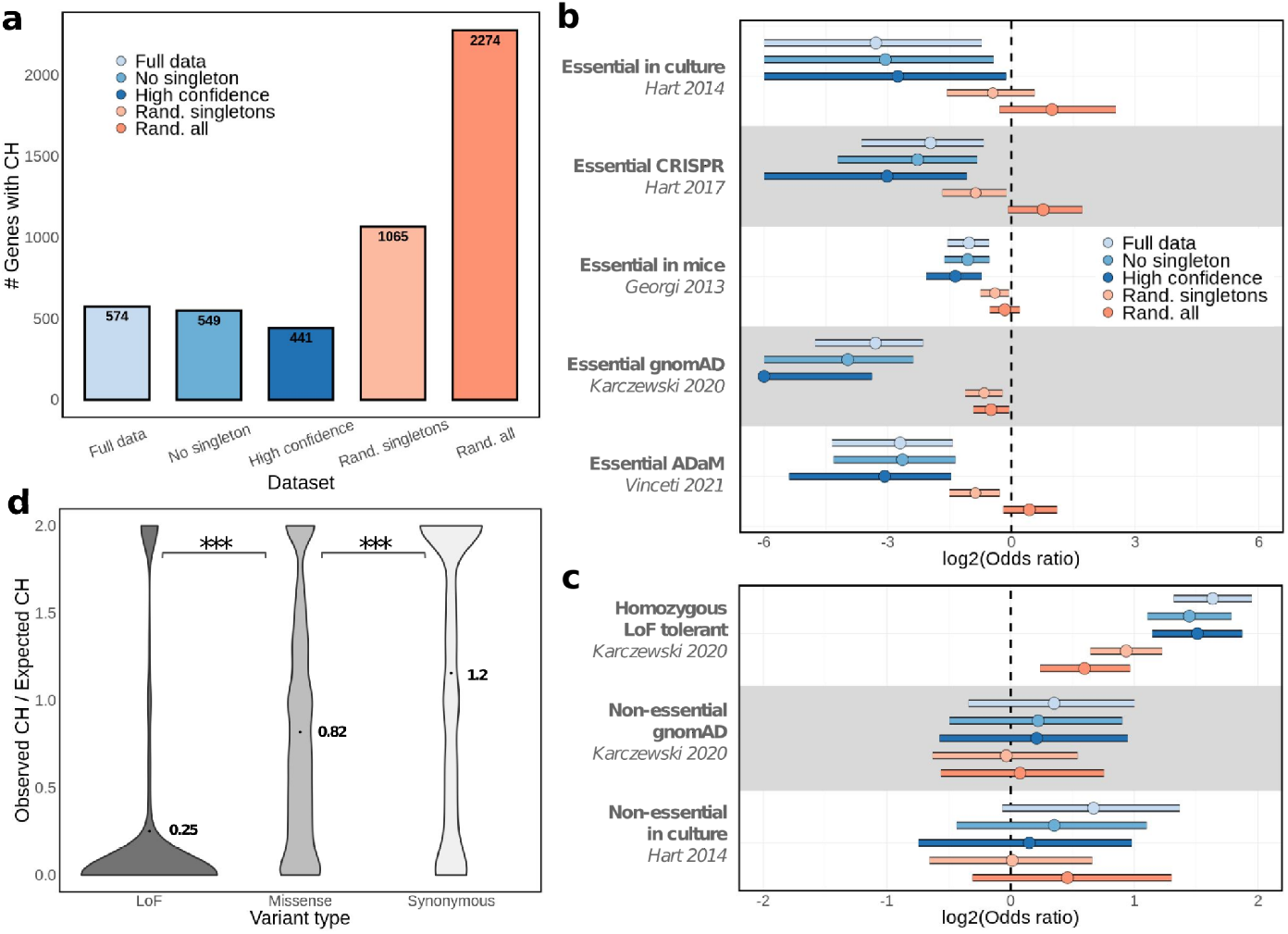
Compound heterozygous identification in the UK Biobank WES data phased with SHAPEIT5. (**a**) Number of genes with at least one individual with compound heterozygous (CH) LoF variants across several categories: “Full data”: all LoF variants in the study, “No singleton”: all LoF variants except singletons, “High confidence”: LoF variants excluding singletons and calls with phasing confidence score < 0.99, “Rand. singletons”: including all LoF variants but shuffling phasing of singletons, “Rand. all”: shuffling phasing of all LoF variants. (**b**) Two-way Fisher’s Exact test odds ratios and 95% confidence interval (log2-scaled) of CH genes versus non-CH genes presence in multiple lists of essential genes (see Methods). Background is composed of 3018 genes with ≥2 LoF mutations. X-axis is capped at -6. (**c**) Same as previous but across lists of non-essential or LoF tolerant genes. (**d**) Ratio between the number of individuals with CH events and the expected number of individuals given the number of variants, per gene. Missense and synonymous variants are shown in addition to LoF variants as a comparison. Values and points refer to the mean ratio. “***” corresponds to Wilcoxon test p-values < 2e^-226^.

As a major feature of SHAPEIT5 is the ability to phase singletons (42% of LoF variants used, **Supplementary Fig. 10**), we evaluated the benefit of singleton phasing in CH identification by comparing to experiments either excluding singletons or phasing singletons randomly, mimicking standard statistical phasing approaches. When singleton variants are excluded from analysis, 549 CH genes and 779 CH events are identified, resulting in a loss of 25 CH genes and 50 CH events compared to the full dataset (**Figure 3a**). Excluding CH events detected with singletons does not significantly change the enrichment/depletion of essential/non-essential gene lists, indicating that the CH genes identified with singletons are not significantly different from other CH genes (**Figure 3b**,**c**). When singletons are assigned to one of the two haplotypes randomly (i.e. random phasing), 1,065 CH genes and 1,370 CH events are identified (**Figure 3a, Supplementary Fig. 11**). However, the additional genes identified reduce the depletions in essential genes previously observed (**Figure 3b**), which confirms the need for accurate phasing to determine CH events. The results obtained when randomizing singletons are similar to those obtained when phasing with Beagle5.4, on which 1,207 CH genes (1586 CH events) are identified, but only mild or non-significant depletions to essential genes are observed (**Supplementary Fig. 12**). When excluding singletons, Beagle5.4 identifies 673 CH genes (962 CH events) which are significantly depleted in essential genes but at reduced levels compared with SHAPEIT5 phasing (**Supplementary Fig. 12**). Finally, as a control, we randomized both singletons and other variants, which led to 2,274 CH genes and 17,886 CH events (**Figure 3a**), which did not display depletion in essential genes, as expected (**Figure 3b**). Together, these results indicate that accurate haplotype phasing is crucial for the identification of CH events and that including singletons phased with SHAPEIT5 leads to *bona fide* discoveries.

The finding that CH genes are depleted in essential genes indicates that CH events are avoided, at least in a subset of the genes. To explore this further, we compared the number of expected and observed CH events per gene, based on the variant distribution in the UK Biobank cohort, assuming that each variant phase is independent (see Methods). In the average gene, we found that only 25% of the expected number of individuals with CH are observed, confirming evidence for negative selection (**Figure 3d**). Importantly, when performing CH gene and event discovery using variants with synonymous effect (**Supplementary Fig. 13, Supplementary Data 1**), the number of observed CH generally matches the expectation (mean ratio = 1.2, **Figure 3d**), indicating no or low selective pressure to reduce synonymous variant CH events for most genes. When considering missense or low-confidence LoF variants (referred to as *missense* for simplicity), we observed a mild decrease in observed CH events compared to expected (mean ratio = 0.82, **Figure 3d, Supplementary Data 1**), consistent with the possible deleterious effect of some missense variants. In addition, we found that missense CH genes had only mild or no depletion for essential genes compared to LoF CH genes, whereas synonymous CH genes either had no significant depletions or were even enriched in some essential gene sets (**Supplementary Fig. 13**). Overall, our results demonstrate that the accurate phasing at rare variants with SHAPEIT5 allows us to screen for CH events across the UK Biobank cohort with high confidence, revealing that LoF CH events are under strong selective pressure in essential genes, as expected by their high negative impact.

## Discussion

In this work we described a new method, SHAPEIT5, to accurately phase millions of rare variants from large sequencing datasets. We applied this method on the UK Biobank to produce phased genomes for the available whole genome and exome sequencing data for a total compute cost below 4,000 British pounds. The resulting haplotype estimates have low switch error rates, with rare variants down to doubletons being phased with high confidence, which can be leveraged for a high genotype imputation accuracy at rare variants when used as a reference panel. Beyond measuring error rates as commonly done, we also biologically validated phased haplotypes by identifying CH events, which we found highly depleted in essential genes, as expected. In addition, we also achieved singleton phasing, albeit with higher error rates. When analyzed collectively, the phased singletons provided additional reliable discoveries of compound heterozygous events.

Although of significant interest, previous knowledge of compound heterozygous cases come mostly from study cases in families^7,8^ and there is currently no existing method to systematically identify CH events in large biobanks. Here, we show that high-quality phasing of rare variants with SHAPEIT5 allows studying compound heterozygosity at the biobank-scale level, which can greatly increase the number of CH events characterized compared to the use of family data, in addition to exploring their association with novel phenotypes. As a proof-of-principle, we screened all protein-coding genes for CH events with high-confidence loss-of-function variants and found 549 genes fully knocked out across 816 UK Biobank individuals out of the 374,826 individuals considered in this study. This corresponds to 0.22% of the population having at least one gene knockout by compound LoF heterozygous events. The frequency of CH events observed matches previous estimates on the amount of CH events likely to be observed in outbred healthy cohorts of ∼400.000 individuals^30^. We thus provide a key resource that complements other lists of non-essential genes^31^, with the main advantage that these knockouts are found *in vivo* in generally healthy participants. Indeed, UK Biobank participants are not expected to have any rare and/or severe genetic diseases as their average age is 56 years (CH individuals mean age = 56.5), which is after the age of onset for most rare diseases. Truly deleterious CH events are expected to be depleted in this cohort compared to the general population which partially explains that the gene knockouts we observed are strongly depleted in multiple lists of essential genes. However, we still found 52 genes deemed as essential in at least one of the essential gene lists we analyzed. We can conceive three possible scenarios to explain these specific cases. First, the CH event had a moderate impact on the individual and did not result in severe disease. As an example, we found one individual with pulmonary embolism while having the essential gene *ADAM19* knocked-out, a gene reported for its involvement in pulmonary disease^32,33^. Second, compensatory mutations can rescue the deleterious effect of the knockout. For instance, we found one individual having a knockout of CFTR, an essential gene found to be rescued by multiple gain-of-function mutations across the genome^34–36^. Finally, some of the discovered CH events may be false positives driven by incorrect phasing or erroneous LoF annotations.

We foresee that rare variant phasing in large sequencing studies such as the UK Biobank has the potential to unlock many applications and analyses. First, other types of functional variants can be screened for compound heterozygous effects, for instance, combining LoF and missense or regulatory variants^37^. Second, phase information can be included in rare variant burden testing approaches, which usually consider only a mixture of the two haplotypes. Third, using accurately phased reference panels allows to phase extremely rare variants with high accuracy, even singletons to some extent, for any new sequenced genome from the same population. This is beneficial for diagnosis of rare and severe diseases caused by compound heterozygous effects, such as in the Genomics England dataset^38^, in which diagnosis yield could potentially be increased by incorporating phase information.

## Methods

### Common variant phasing

For common variant phasing (MAF ≥ 0.1%), SHAPEIT5 is largely based on the previous SHAPEIT version (i.e. version 4). Briefly, it updates the phase of each sample in turn within a Gibbs sampler iteration scheme: each sample is phased by conditioning on other samples’ haplotypes using the Li and Stephens model^22^. Two main improvements, part of the SHAPEIT4 model, allow fast phasing at common variants. First, the haplotype sampling step has linear complexity in the number of conditioning states^39^ and is multi-threaded so that multiple samples are phased in parallel. Second, the sampling is based on a parsimonious and highly informative set of haplotypes, identified in constant time using the Positional Burrows Wheeler Transform data structure. However, one computational limitation of SHAPEIT4 resides in its inability to parallelize the construction of the PBWT, which can become relatively long in very large datasets. In SHAPEIT5, we introduced a parallelization scheme for the PBWT construction: multiple PBWT passes are run in parallel on multiple CPU cores, each one running for a different chunk of 4cM by default, achieving a significant reduction of the wall clock running time of the method.

### Rare variant phasing

In order to accurately phase rare variants (MAF < 0.1%), SHAPEIT5 uses the haplotypes derived at common variants as haplotype scaffolds onto which heterozygous genotypes are phased one rare variant at a time. For a single heterozygous genotype we aim to determine which of the two target’s chromosomes carries the minor allele (as opposed to the major allele). To do so, our method uses the Li and Stephens model to compute the probabilities of the two possible phases. The probabilistic inference is based on a set of haplotypes carried by other samples in the dataset, that we call conditioning haplotypes (or samples depending on the context). The outcome of the estimation is a posterior probability of the most likely phase for each of the rare heterozygotes. Specifically, our model comprises five main features:

#### Sparse representation

We use a sparse matrix representation of the genotypes at rare variants to efficiently store large amounts of genotype data in memory and speed up computations. Only genotypes carrying at least one copy of the rare allele are stored in memory together with the necessary indexes to determine the sample and variant to which the genotype corresponds. As most of the rare variants are homozygous for the major allele, this representation allows for a large reduction in memory usage and a fast identification of heterozygous genotypes at a given rare variant. To quickly retrieve rare genotypes at both the sample and variant levels, we store this sparse genotype matrix in memory together with its transpose.

#### Haplotype selection

To get the most informative haplotypes in the conditioning set, we require that they (i) share long haplotype matches with the target and (ii) are not monomorphic at the rare variant of interest. The first condition ensures that the haplotypes in the conditioning set are informative for the copying model. The second condition ensures that the conditioning set contains carriers of the two possible alleles at the rare variant of interest. The latter is required to accurately contrast the two possible phasing possibilities of the rare heterozygous variant. To efficiently retrieve haplotypes complying with these properties, we use the Positional Burrows Wheeler Transform (PBWT) data structure of the haplotype data derived at common variants. We perform both forward and backward PBWT sweeps so that we can identify long matches between haplotypes centered in the position of the rare variant by interrogating the flanking prefix arrays. This gives a first set of haplotypes that complies with condition (i), but not necessarily with condition (ii). Therefore, we do a second identification of matches in the PBWT, this time restricting the search to the subset of samples carrying the minor allele. We efficiently achieve this second pass by taking advantage of the sparse genotype representation: we only interrogate the PBWT prefix arrays at the sparse indexes.

#### Forcing homozygosity

The conditioning set defined before contains a set of haplotypes that share large segments with the target haplotype at common variants, but they have not been phased yet at the rare variant of interest. When the conditioning sample is homozygous, this is not an issue as its two haplotypes carry the same allele. However, when the conditioning sample is heterozygous, we do not know the allele carried by each one of its two haplotypes. We solve this by simply assigning the minor allele to both haplotypes. As a consequence of the two previous steps, the conditioning set of haplotypes is guaranteed to contain carriers of the two possible alleles at the rare variant of interest.

#### Copying model

We can now perform phasing of rare heterozygous genotypes based on the conditioning set of haplotypes that have been constructed as part of all the previous steps. SHAPEIT5 computes the probability that each target haplotype carries the minor allele by using a haploid version of the Li and Stephens model. Specifically, it runs a forward-backward pass as done in the context of genotype imputation (see ^40^ for details) to get the probabilities that each target haplotype carries the minor allele at the rare variant. The conditioning set of haplotypes serves as a local reference panel for imputing the alleles at the rare variant in the target sample. Of note, accurate inference is made possible since the conditioning set we chose is guaranteed to comprise carriers of both the major and the minor alleles at the rare variant of interest. Having only carriers of a single allele would not be informative for making inference here. Finally, we use these imputation probabilities to derive phasing probabilities (**Supplementary Fig. 1**), which we can use to get the most likely phase or to propagate phasing uncertainty in downstream analyses.

#### Singleton phasing

In the case of singletons, only the target sample carries a copy of the minor allele at the rare variant. Therefore, none of the conditioning haplotypes carries the minor allele and the whole copying model described above is unable to make inference. This is a well-known limitation of all statistical phasing methods. SHAPEIT5 can provide inference at these sites by using the Viterbi algorithm for the Li and Stephens model ^22^, to obtain the longest shared IBD segment between each one of the two target haplotypes and the conditioning haplotypes. The minor allele of singletons is then assigned to the target haplotype with the shortest shared segment. The idea behind this model presumes that the shorter the IBD sharing between two haplotypes, the older their most recent common ancestor is, and therefore, the chance for new mutations to occur is increased in that lineage.

### Validation of haplotype estimates

To validate haplotype estimates, we use trios (two parents, one offspring) for WES data and both duos (parent-offspring pairs) and trios for WGS data. To identify parent-offspring relationships, we use the kinship estimate and the IBS0 as provided as part of the UK Biobank SNP array release. We select parent-offspring relationships as having a kinship coefficient lower than 0.3553 and greater than 0.1767 and an IBS0 lower than 0.0012^1,41^. In addition, we require that the difference in age between parents and offspring is greater than 15 years and that the two parents have different sex for trios. We finally keep only self-declared white British individuals for which ancestry was confirmed by Principal Component Analysis (PCA, UKB field 22006). The number of trios used in the validation for all three datasets (Array, WES or WGS) is shown in **Supplementary Tables 1**. Validation of haplotypes is a two-step procedure. First, we statistically phase a given dataset including only the offspring samples. Second, we use the parents to measure the switch error rate (SER), a commonly used metric to assess how close estimated and true haplotypes are. The SER is defined as the fraction of successive pairs of heterozygous genotypes being correctly phased. In the context of this work, we measured SER stratified by bins of minor allele count (MAC). We assigned each heterozygous genotype to a given MAC bin and counted the fraction of heterozygous genotypes being correctly phased per MAC bin. This definition of SER has the advantage of showing how well statistical phasing performs depending on the frequency of the variants it phases (either common or rare).

### UK Biobank SNP array dataset

We used the UK biobank Axiom array in PLINK format and converted it into VCF format using plink2. This resulted in 784,256 variant sites across autosomes for 488,377 individuals. We then applied quality control on the data using the UK biobank SNPs and samples QC file (UK Biobank Resource 531) to only retain SNPs and individuals that have been used for the official phasing of the Axiom array data^1^, resulting in 670,741 variant sites across 486,442 individuals. This includes 897 white British parent-offspring trios and 4373 white British parent-offspring duos (**Supplementary Table 1**).

### UK Biobank WGS dataset

We use the Whole genome GraphTyper joint call pVCFs from the UK Biobank RAP. We first decomposed multi-allelic variants into bi-allelic variants using bcftools norm -m^42^. We then performed quality control of the variant sites and filtered out SNPs and indels for (i) Hardy-Weinberg p-value < 10^−30^, (ii) more than 10% of the individuals having no data (GQ score=0; missing data), (iii) heterozygous excess less than 0.5 or greater than 1.5, and (iv) alternative alleles with AAscore < 0.5. Additionally, we kept only variant sites with the tag “FILTER=PASS”, as suggested by the data providers^43^. This resulted in a total of 603,925,301 variant sites, including 20,662,402 common variant sites (MAF ≥ 0.1%) and 583,262,899 rare variant sites (MAF < 0.1%), across a total of 150,119 individuals. This WGS dataset includes 31 trios and 432 duos (**Supplementary Table 1**). To assess the accuracy of the phasing, we use chromosome 20 only. For this analysis, we only used samples being also genotyped with the UK Biobank Axiom array, resulting in 147,754 individuals (**Supplementary Table 1)**. We phased chromosome 20 using chunks of on average 4.5Mb with overlapping buffers of 250kb. We used Beagle5.4^15^ with default parameters on the entire chromosome 20.

### UK Biobank WES dataset

We used the WES files in pVCF format as released on UK Biobank Research Analysis Platform (RAP). Quality control pipeline has been described in Szustakowski et al.^44^. To phase WES data, we first merged it with the SNP array data to increase the number of common variants and therefore improve the quality of the haplotype scaffold onto which rare variants are phased. We kept only individuals with both the SNP array and the WES data, resulting in 452,644 total individuals, including 719 white British parent-offspring trios and 3014 white British parent-offspring duos. When a variant is listed in both the WES and the SNP array, we keep the SNP array copy as the SNP array is expected to be more robust to SNP calling errors^14^. This resulted in retaining a total of 26,199,614 variants, including 977,517 common variants (MAF ≥ 0.1%) and 25,222,097 rare variants (MAF < 0.1%) (**Supplementary Table 1**). Phasing the 452,644 individuals with both WES and Axiom array available data is performed for each chromosome independently in a single chunk. We also used Beagle5.4^15^ with default parameters.

### Genotype imputation

To perform genotype imputation from the phased WGS and WES datasets, we extracted 1,000 samples with British ancestry that are unrelated to any other sample in the dataset, and for which we had Axiom SNP array data available. We therefore used a reference panel composed of the remaining 146,754 WGS samples and 446,470 WES samples for both SHAPEIT5 and Beagle5.4. For the HRC reference panel, we used PICARD toolkit (URL: http://broadinstitute.github.io/picard/) to liftover the data to the Human genome assembly GRCh38, retaining 99.8% of the original variants.

We used Beagle5.4^15^ for genotype imputation of SNP array data, allowing pre-phasing from the reference panel. We accessed imputation accuracy by measuring the non-reference discordance rate and the squared Pearson correlation between imputed and high-coverage genotypes using the GLIMPSE_concordance tool ^45^ (-gt-val option) at custom allele count bins (--ac-bins 1 5 10 20 50 100 200 500 1000 2000 5000 10000 20000 50000 100000 146754 for WGS, --ac-bins 1 5 10 20 50 100 200 500 1000 2000 5000 10000 20000 50000 100000 446470 for WES). A drop of correlation quantifies the reduction in effective sample size in association testing due to imperfect imputation. For instance, a difference of 0.05 involves a power loss equivalent to losing 5% of the data.

We also evaluated the non-reference discordance rate using the GLIMPSE_concordance tool. The non-reference discordance is calculated as NRD = (*e*_*rr*_+*e*_*ra*_+*e*_*aa*_)/(*m*_*ra*_+*m*_*aa*_+*e*_*rr*_+*e*_*ra*_+*e*_*aa*_), where e_rr_, e_ra_ and e_aa_ are the counts of the mismatches for the homozygous reference, heterozygous and homozygous alternative genotypes respectively, and m_ra_ and m_aa_ are the counts of the matches at the heterozygous and homozygous alternative genotypes. NRD is an error rate that excludes the homozygous reference matches, which are the majority at rare variants, giving more weight to the other matches. We computed the non-reference discordance rate within frequency bins in the reference panel.

### Compound heterozygosity detection

We restricted the analysis to the cohort of self-declared white British individuals for which the ancestry is confirmed by Principal Component Analysis (UKB field 22006) with both SNP array and exome-seq data, excluding parental individuals (N = 374.826). Only WES variants with MAF <0.1% before sample filtering were considered. Variant annotations (LoF, Synonymous and Missense|LC) were obtained from the Genebass database^46^ through Hail (gene-level results, results.mt). Briefly, these variants had been annotated by Ensembl VEP v95^47^ and LoF variants (stop-gain, frameshift and splice donor/acceptor sites) were further processed by LOFTEE^4^, separating high-confidence (used as “LoF”) from low-confidence (used in the “Missense|LC’’ category). Only unique canonical transcripts for protein-coding genes were considered. LoF, synonymous and missense variants were gathered in the UKB cohort using bcftools isec function, with “-c none” parameter to match variants by chromosome, position, reference and alternative alleles.

Identification of compound heterozygous events was performed with custom Python (v3.7) scripts. Briefly, for each variant type (LoF, synonymous, missense) and for each gene, individuals with at least 2 mutations were assessed for compound heterozygosity by having at least 1 variant in each of the two haplotypes. In addition, for each gene, we calculated the expected number of individuals with CH as 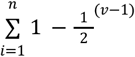, where *v* indicates the number of variants in individual *i*. To compare the number of LoF CH genes and events without phasing we randomized (i) phasing at singletons (defined as MAC = 1 when considering all participants with WES data) and (ii) phasing at all variants, by attributing 0.5 probability for each variant to fall in either of the two haplotypes, independently for each variant.

### Essential and non-essential gene lists

We obtained lists of essential and non-essential genes from multiple sources (described below). For each of these gene lists, we performed Fisher’s Exact tests (two-sided) for several categories of CH genes versus non-CH genes, considering a background of 3,018 genes with at least one individual with two LoF mutations. For synonymous and missense variants, the background included 10,514 and 15,612 genes, respectively.

1. “Essential in mice” (N = 2,454) from Georgi et al. 2013^48^ includes genes where homozygous knockout in mice results in pre-, peri- or post-natal lethality and was extracted with ortholog human gene symbols from McArthur’s laboratory^49^ ;
2. “Essential in culture” (N = 360) core essential genes from genomic perturbation screens were obtained from Hart et al. 2014^31^;
3. “Non-essential in culture” (N = 927) putatively nonessential genes (shRNA screening) were obtained from Hart et al. 2014^31^;
4. “Essential CRISPR” (N = 684) genes essential in culture from CRISPR screening were obtained from Hart et al. 2017^50^;
5. “Essential ADaM” (N = 1,075) genes annotated by the ADaM analysis of a large collection of gene dependency profiles (CRISPR-Cas9 screens) across 855 human cancer cell lines (Project Score and Project Achilles 20Q2) were obtained from Vinceti et al. 2021^51^;
6. “Essential gnomAD” (N = 1,920) genes at the bottom LOEUF decile from gnomAD v2.1.1 (i.e. most constrained genes) obtained from https://gnomad.broadinstitute.org/^4^;
7. “Non-essential gnomAD” (N = 1,919) genes at the top LOEUF decile from gnomad AD v2.1.1 (i.e. least constrained genes) obtained from https://gnomad.broadinstitute.org/^4^;
8. “Homozygous LoF tolerant” (N = 1,815) genes with homozygous LoF variants observed in the gnomAD cohort were obtained from Karczewski et al. 2020^4^ (Supplementary Data 7);

## Data availability

The benchmarks on the UK Biobank data have been conducted using the UK Biobank Research Analysis Platform under Application Number 66995. The lists of compound heterozygous events and genes (Supplementary Data 1) are available at (https://docs.google.com/spreadsheets/d/1t_rSZaPCC3ZqrULqNJqPoZ7d9f9wBq3aEd5L8nqE2cc/edit?usp=sharing).

## Code availability

SHAPEIT5 is available under MIT license at https://github.com/odelaneau/shapeit5. This includes code to the phase_common, phase_rare variants, ligate and switch tools and the scripts used to phase WES and WGS data on the UK Biobank RAP. The documentation is available at https://odelaneau.github.io/shapeit5.

## Acknowledgements

Authors were funded by a Swiss National Science Foundation (SNSF) project grant (PP00P3_176977). The funders had no role in study design, data collection and analysis, decision to publish, or preparation of the manuscript.

## Author Contributions

O.D. and S.R. developed the method. R.J.H. performed the phasing experiments.

S.R. performed the imputation experiments. D.M.R performed compound heterozygous analysis. All authors wrote and reviewed the manuscript. O.D. designed and supervised the study.

## Competing Interests

The authors declare no competing interests.

## Supplementary Figures

**Supplementary Figure 1:**
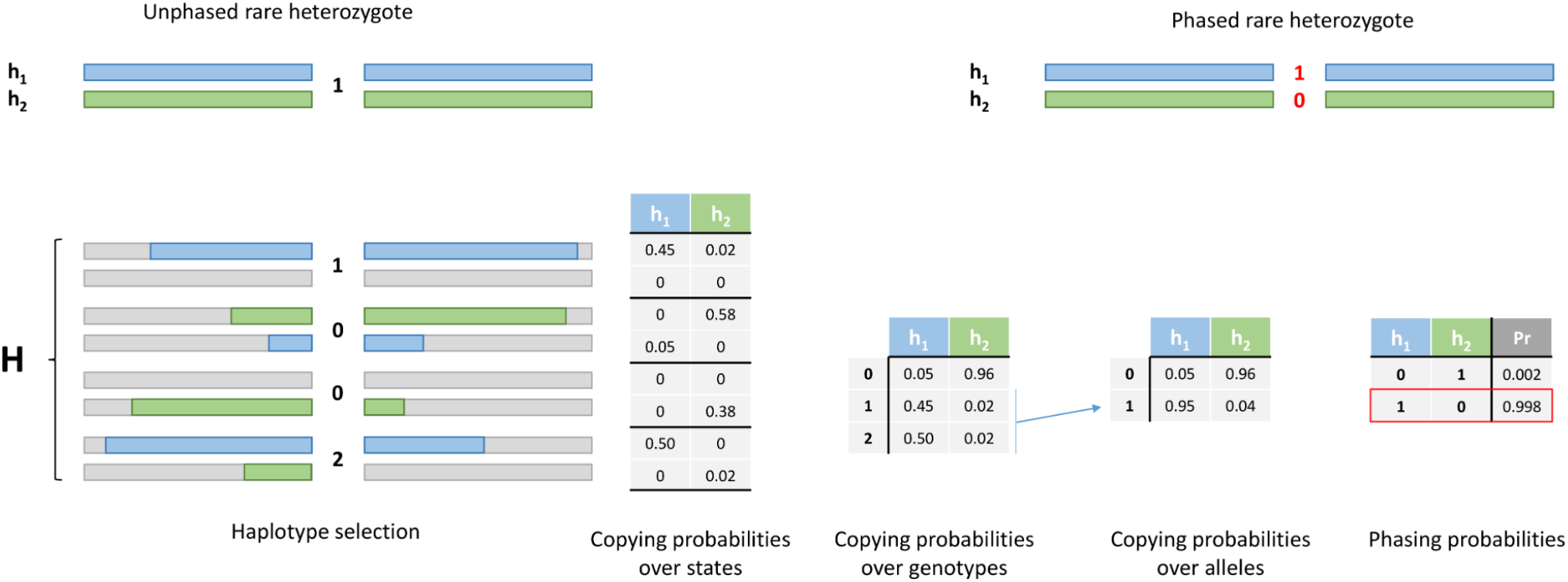
Computation of phasing probabilities. From left to right. A target sample is phased at common variants (h_1_ and h_2_) and is heterozygote for a rare variant (genotype = 1). Haplotypes in the conditioning set H are chosen so that they share long matches with the target (in green and blue). At the rare variant, they are either homozygous for the major allele (genotype = 0), for the minor allele (genotype = 2) or heterozygous (genotype = 1). The haploid Li and Stephens model, as used for genotype imputation, is applied to get copying probabilities, i.e., probabilities that h_1_ and h_2_ copy for each haplotype in H at the rare variant. These probabilities are summed across the possible genotypes at the rare variant. Then, homozygosity is forced at heterozygous genotypes so that the probabilities can be summed per allele (0 or 1) at the rare variant, i.e. the probabilities that h_1_ and h_2_ carry the alleles 0 or 1 at the rare variant. Finally, these imputation probabilities are multiplied in order to get phasing probabilities: (i) h_1_ = 0 and h_2_ = 1 versus (ii) h_1_ = 1 and h_2_ = 0. The most likely phase is then assigned to get the phasing of the rare variant.

**Supplementary Figure 2:**
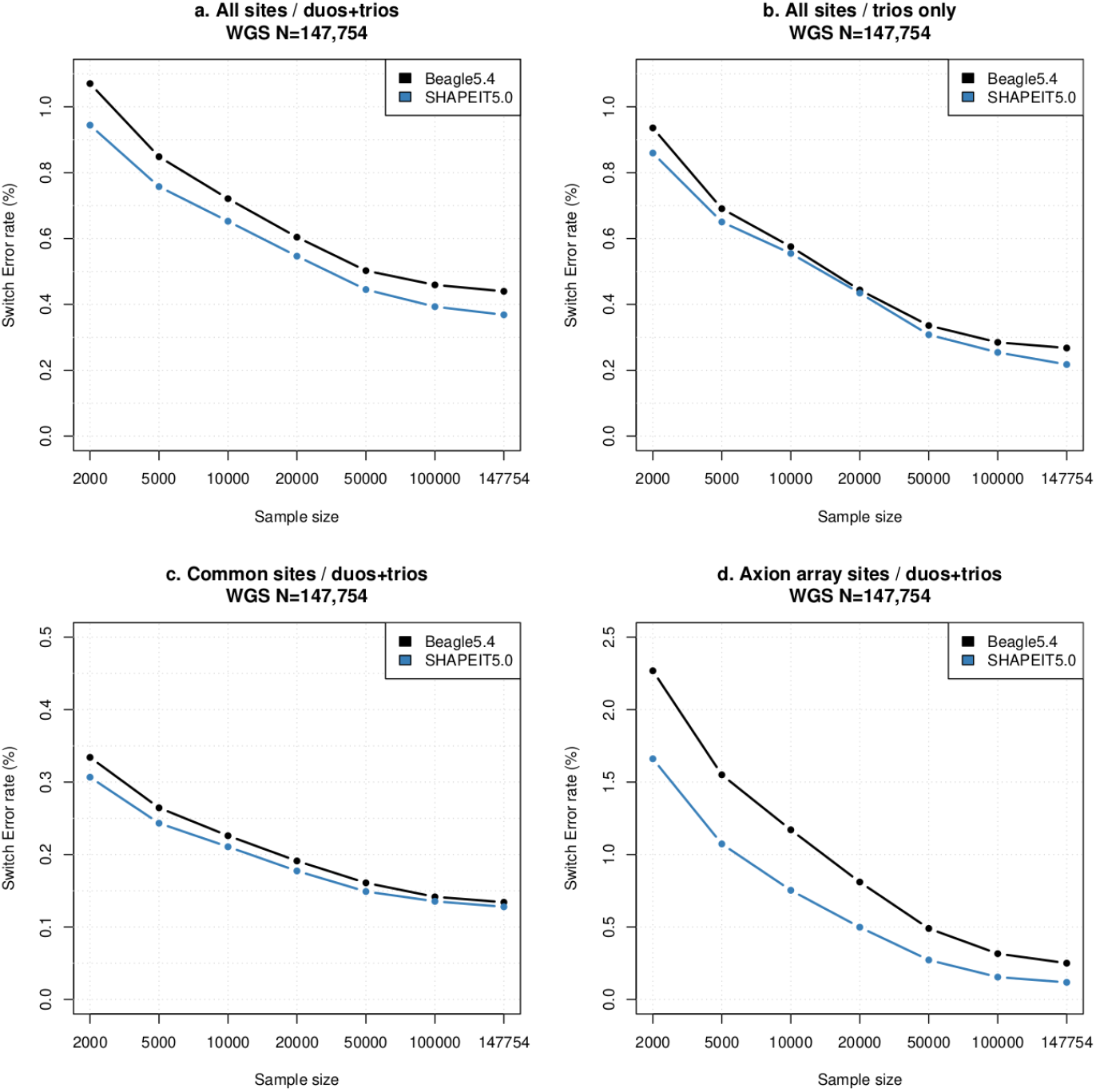
Switch Error rates in the WGS data. (**A**) SER computed at all variants in duos and trios. (**B**) SER computed at all variants just in trios. (**C**) SER computed in duos and trios for common variants (MAF ≥ 0.1%). (**D**) SER computed in duos and trios for variants included on the Axiom array. SER was computed for multiple downsampling experiments comprising 2000, 5000, 10000, 20000, 50000, 100000 and 147754 samples (x-axis). SHAPEIT5 and Beagle5.4 are shown in blue and black, respectively.

**Supplementary Figure 3:**
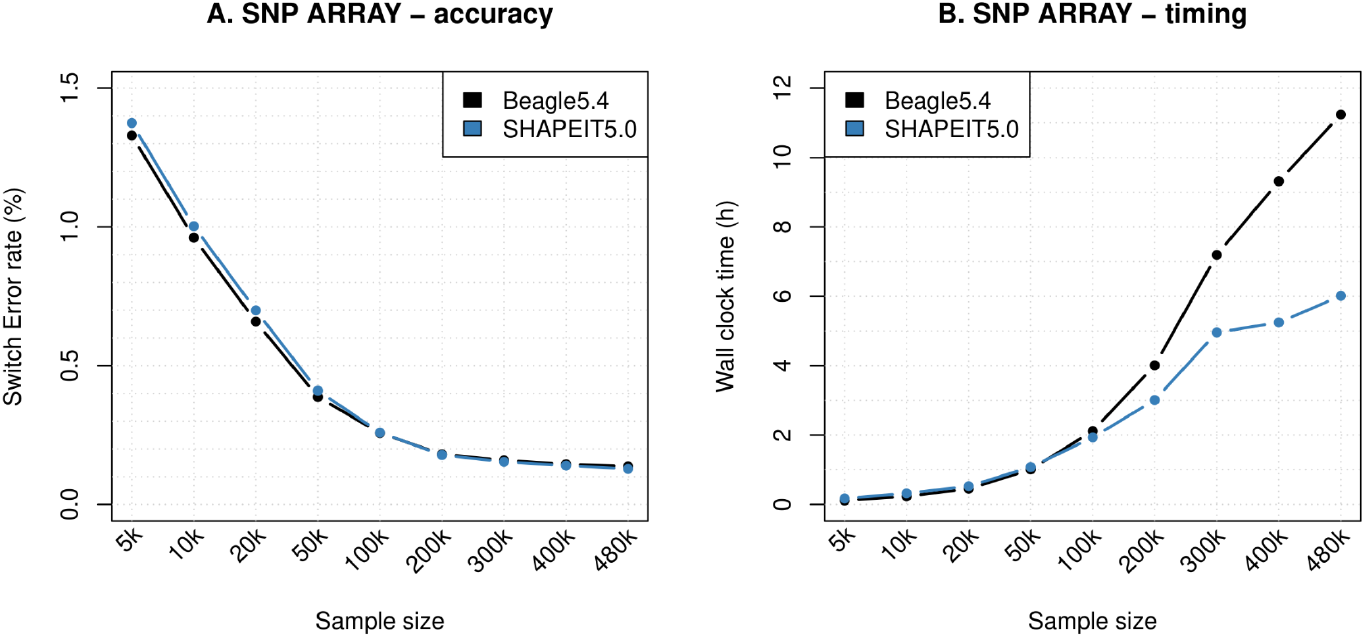
Switch Error rates and running times on the UK Biobank Axiom array data. (**A**) SER computed at all variants typed on the UK Biobank Axiom array in duos and trios. (**B**) Phasing running times on the UK Biobank Axiom array. SER and running times have been computed for multiple downsampling experiments comprising 5000, 10000, 20000, 50000, 100000, 200000, 300000, 400000, 480000 samples (x-axis). SHAPEIT5 and Beagle5.4 are shown in blue and black, respectively.

**Supplementary Figure 4:**
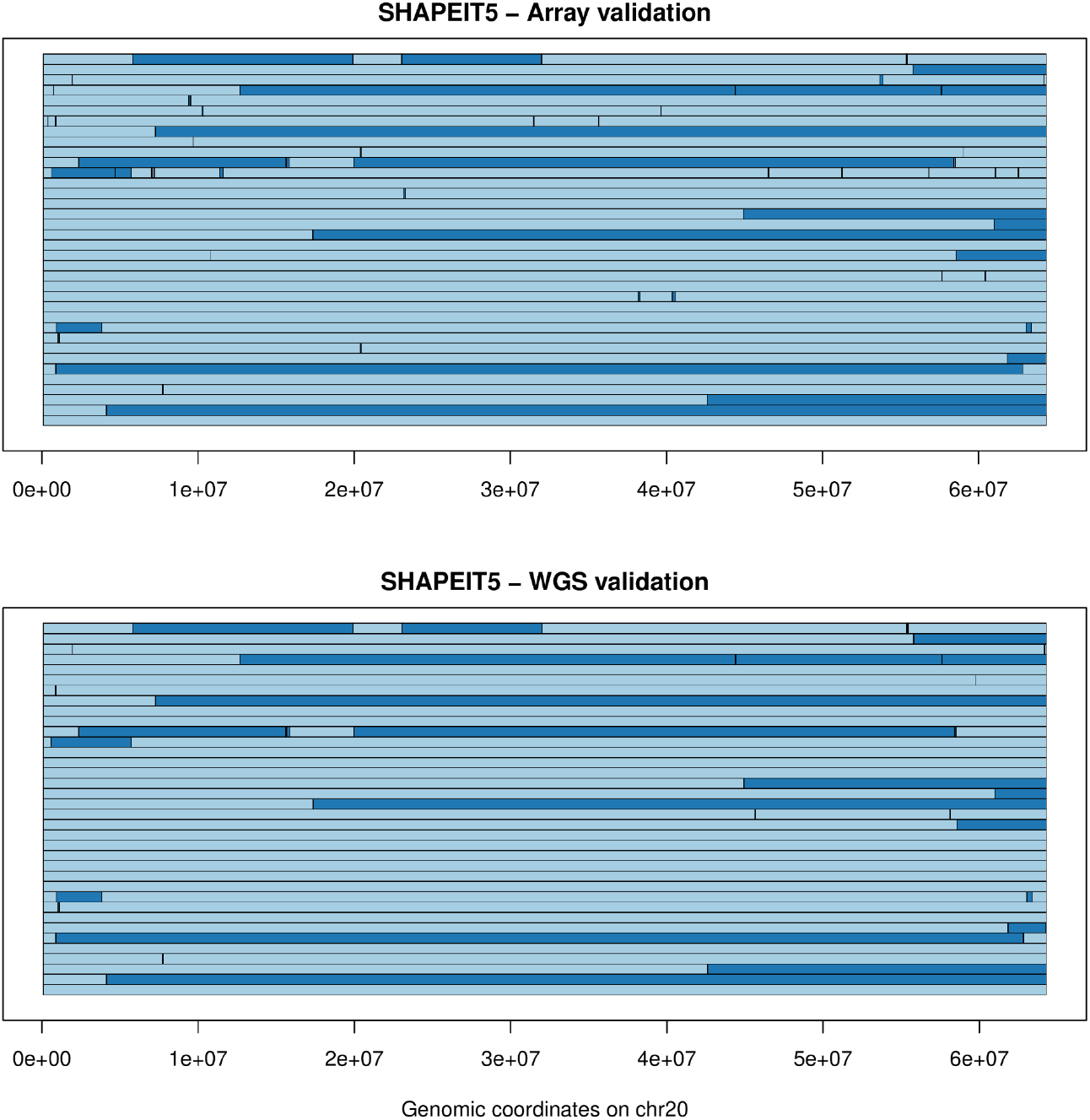
Chromosome 20 switch error locations. Switch error locations for 50 validation samples when phased as part of 480k samples across the entire chromosome 20 (x-axis) with SHAPEIT5. A switch between dark and light blue represents a switch error. At the top is shown the Axiom array phasing when using Axiom array genotypes as validation and at the bottom when using the WGS data as validation genotypes. Many of the small segments in the top panel are due to genotyping errors in the duo/trio parents, which involve incorrect duo/trio phasing as validation. When using the WGS as validation, many of these inconsistencies disappear.

**Supplementary Figure 5:**
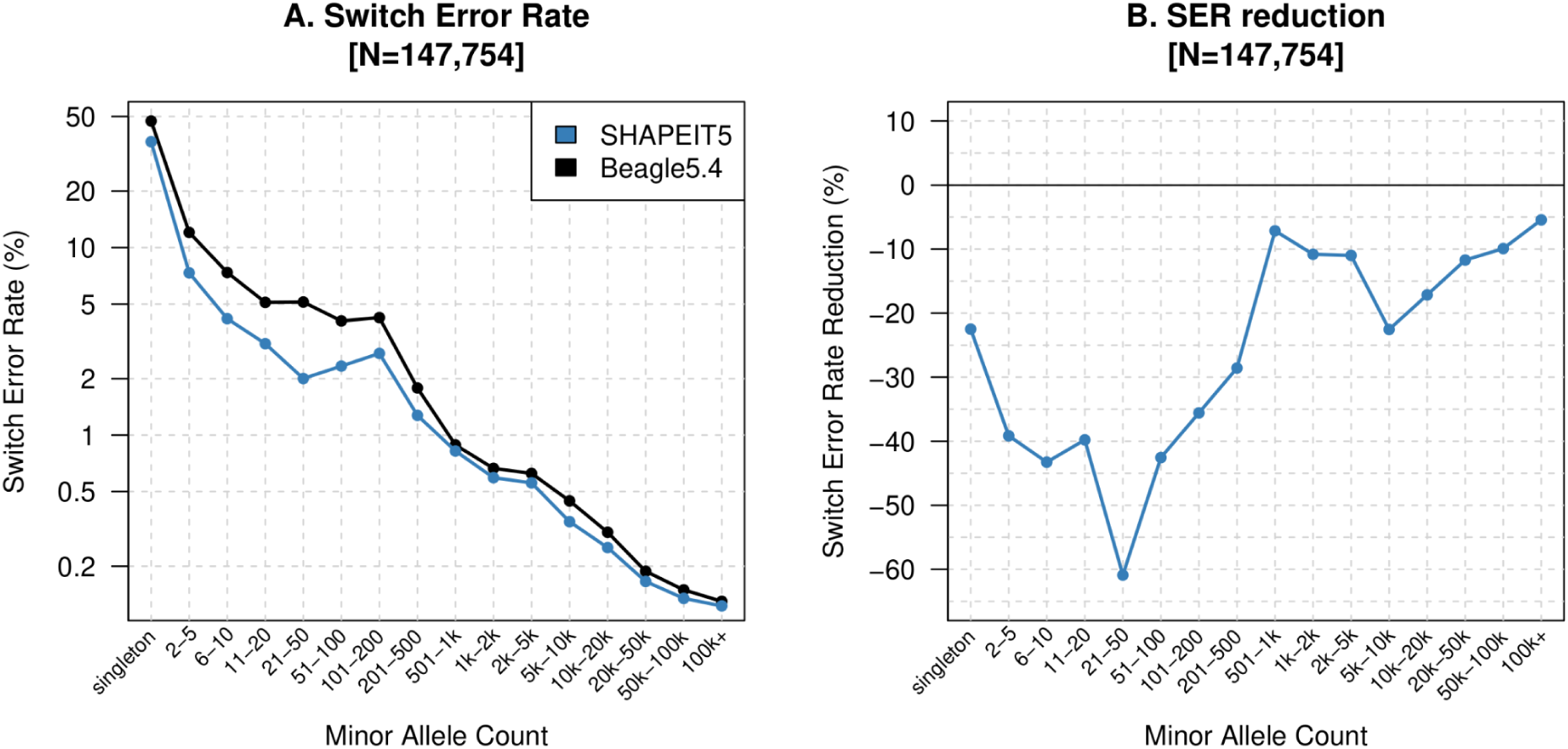
Validation of WGS phasing using trios only. (**A**) SER stratified by minor allele count when using only trios as validation (instead of duos and trios). (**B**) Corresponding SER reductions. Results for SHAPEIT5 and Beagle5.4 are shown in blue and black, respectively.

**Supplementary Figure 6:**
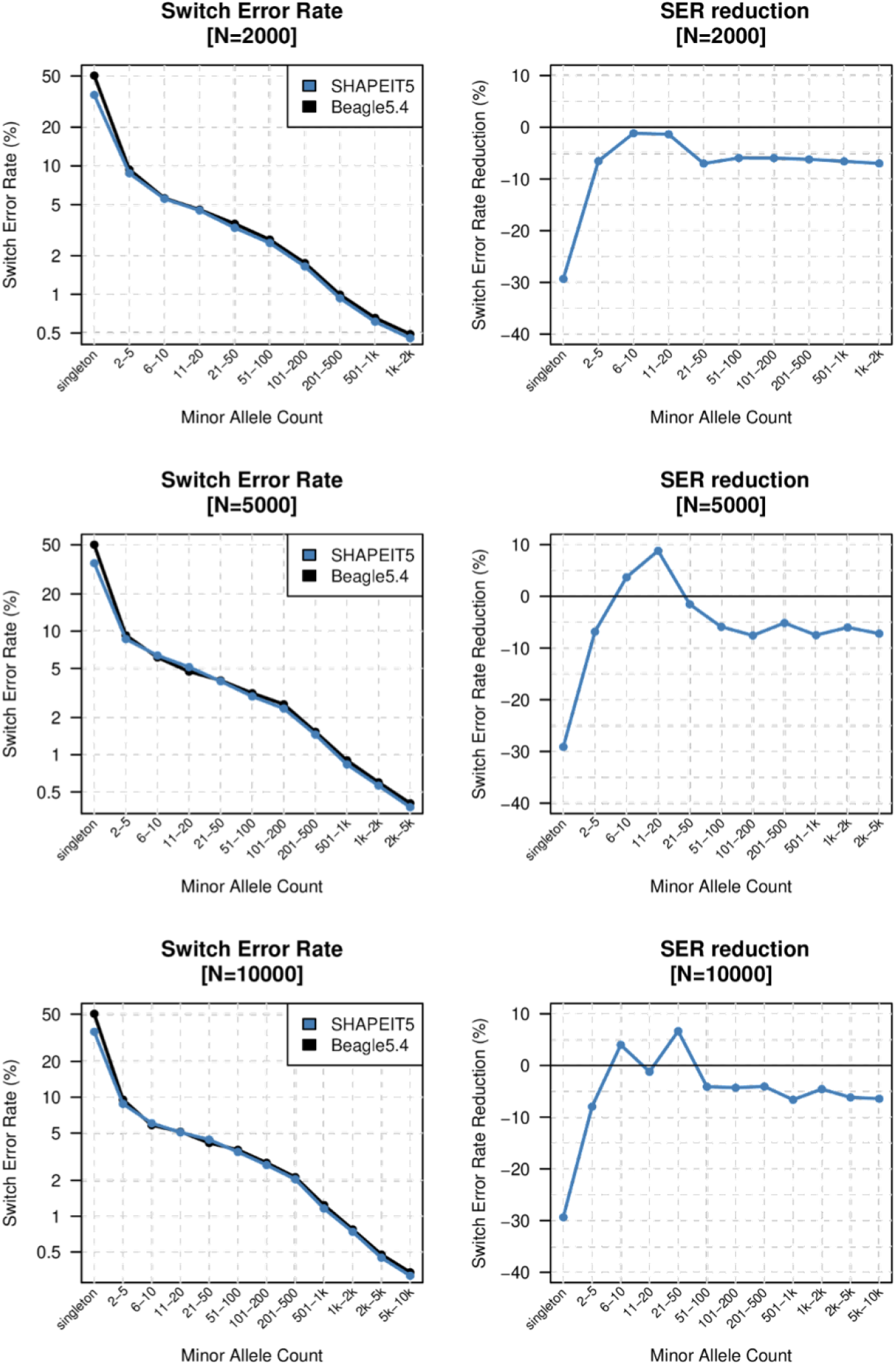
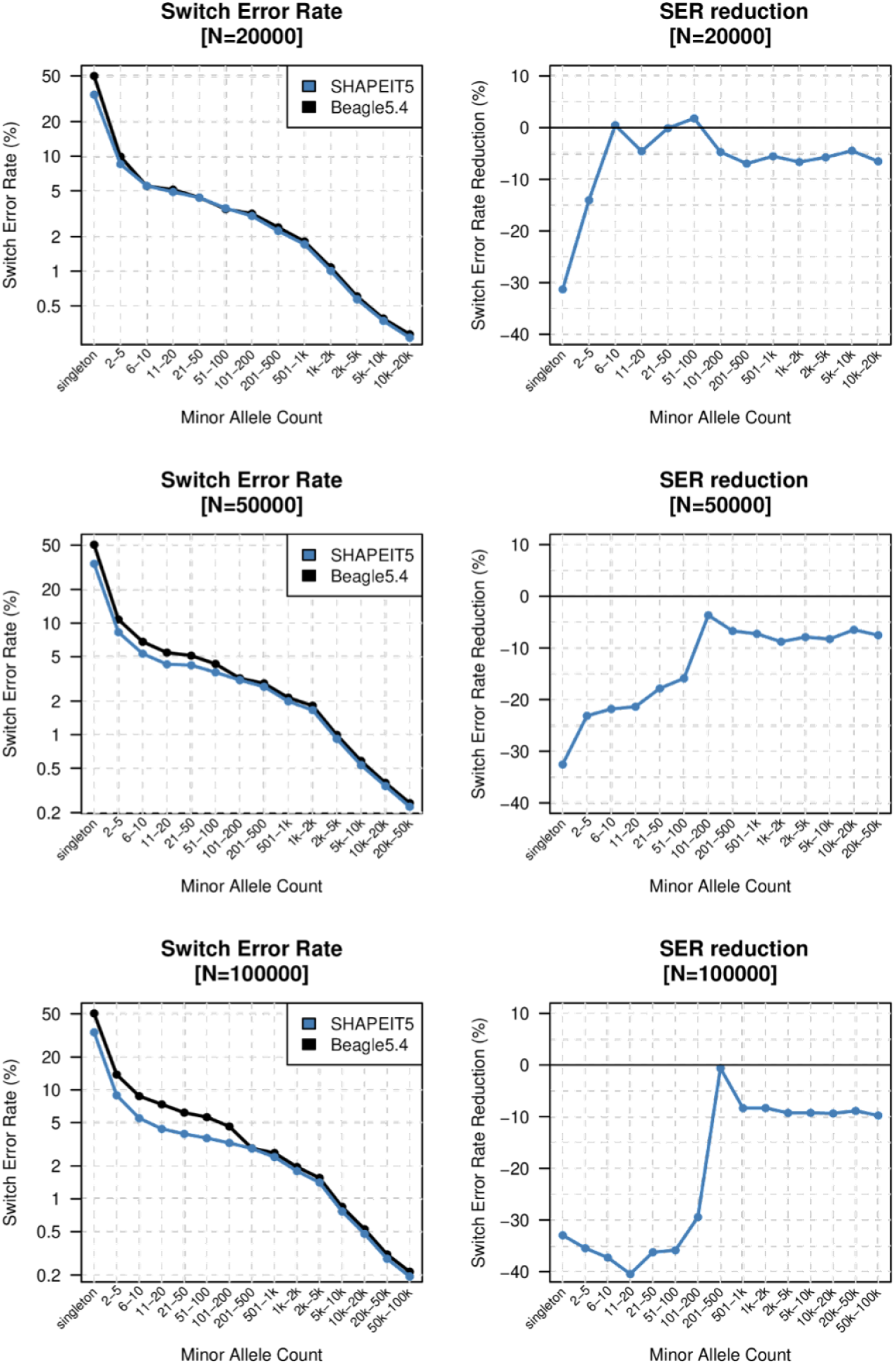
Validation of WGS phasing across multiple sample sizes. Plots on the left show the SER stratified by minor allele count when using trios and duos as validation for, top to bottom, 2000, 5000, 10000, 20000, 50000, 100000 samples. Results for SHAPEIT5 and Beagle5.4 are shown in blue and black, respectively. Plots on the right show the corresponding SER reductions.

**Supplementary Figure 7:**
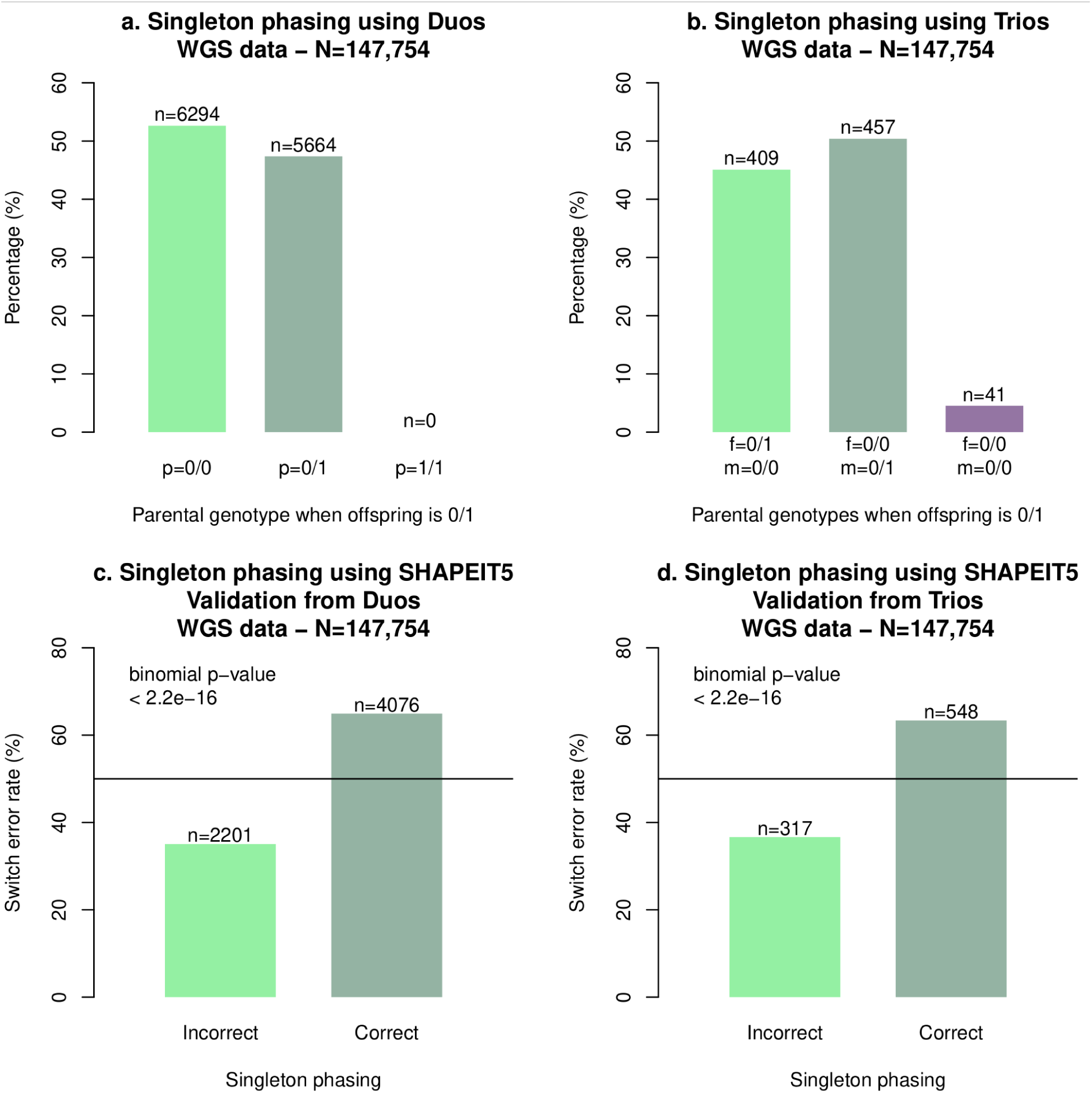
Singleton phasing. (**a**) Numbers of genotyped parents in duos being homozygous for the major allele (0/0) or heterozygous (0/1) for offsprings being heterozygous at the singleton (0/1). Of note, a given variant is assumed to be a singleton here once the genotyped parent is excluded from the dataset to phase. (**b**) Numbers of parents in each genotype class for trios. (**c-d**) Numbers of singletons being incorrectly and correctly phased using duos (**c**) or trios (**d**) as validation.

**Supplementary Figure 8:**
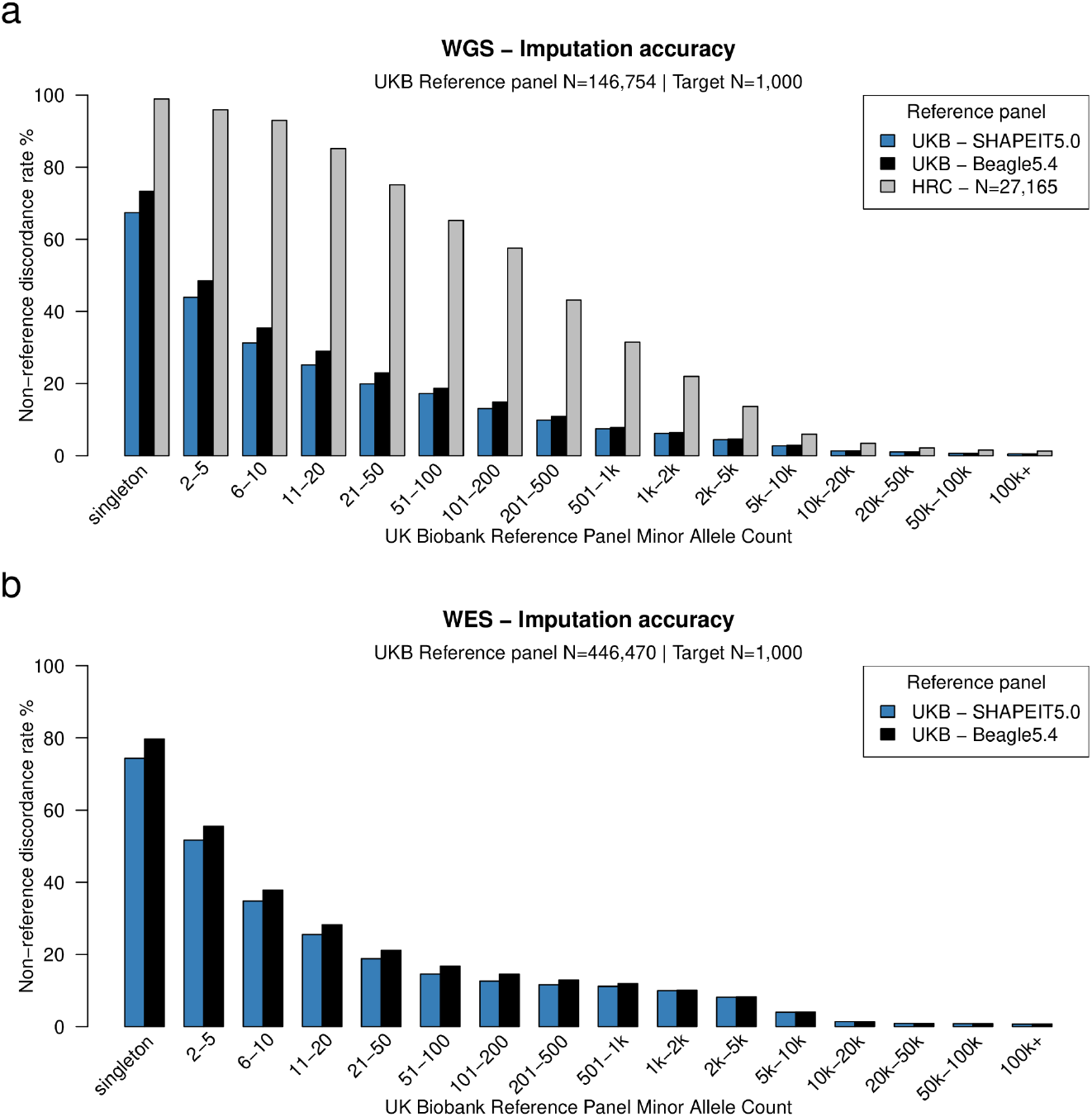
Non-reference discordance rate comparison between SHAPEIT5 and Beagle5.4. Non-reference discordance rate (NRD, y-axis) stratified by minor allele count (x-axis) for 1,000 white British samples genotyped with the Axiom array when using reference panels phased with either SHAPEIT5 (blue) or Beagle5.4 (black) whole genome sequencing (**a**) or whole exome sequencing (**b**). In (**a**) the NRD is also reported for the imputation experiment using the HRC reference panel (gray).

**Supplementary Figure 9:**
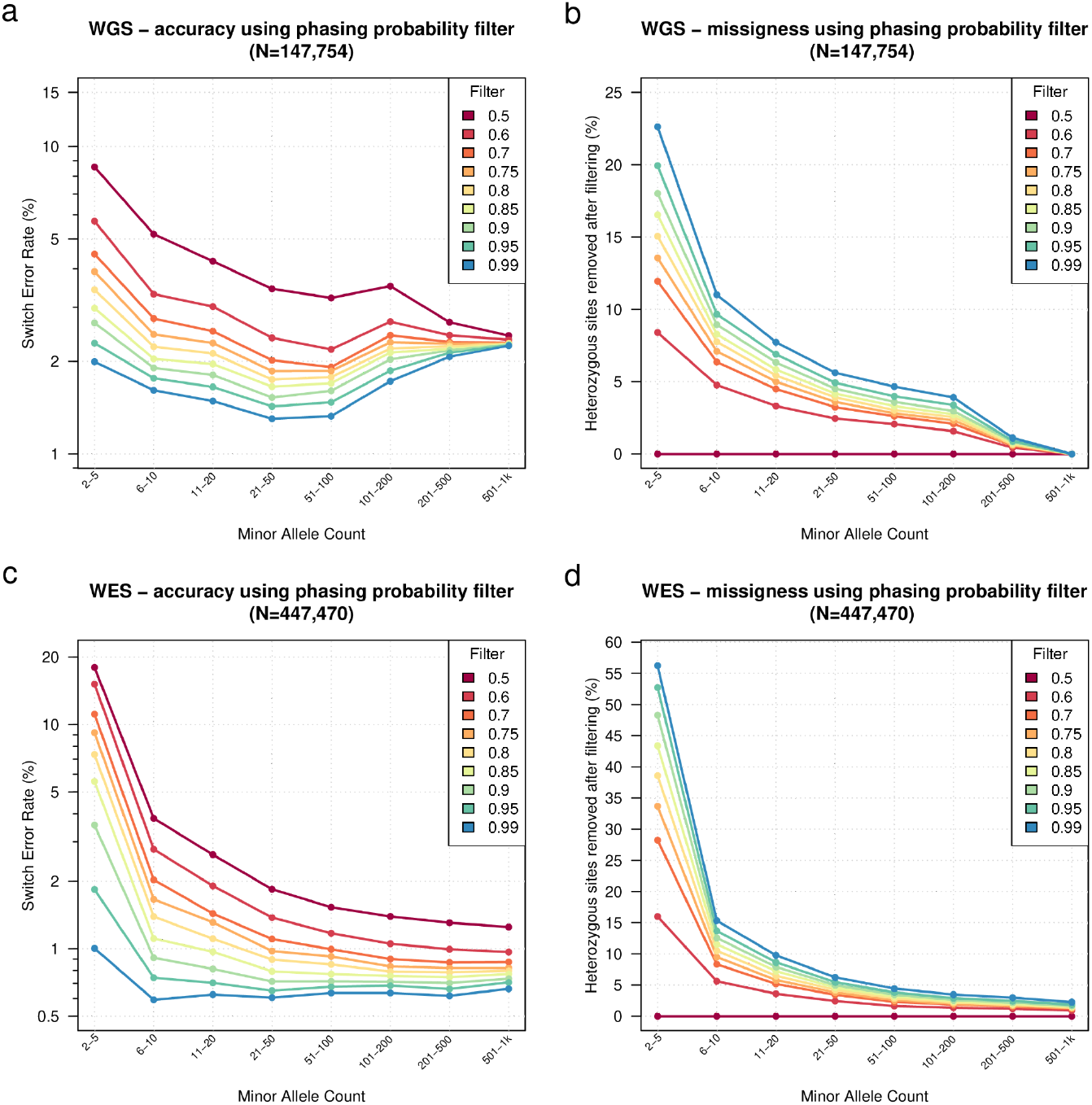
Performance of SHAPEIT5 phasing confidence score. Switch Error Rate (SER, y-axis) for WGS (**a**) and WES (**c**) datasets stratified by minor allele count (x-axis) at different phasing confidence score thresholds. Number of heterozygous sites filtered out at different phasing confidence score thresholds in the WGS (**b**) and WES (**d**) datasets. Filtering for a confidence score of 0.5 is equivalent to no filtering.

**Supplementary Figure 10:**
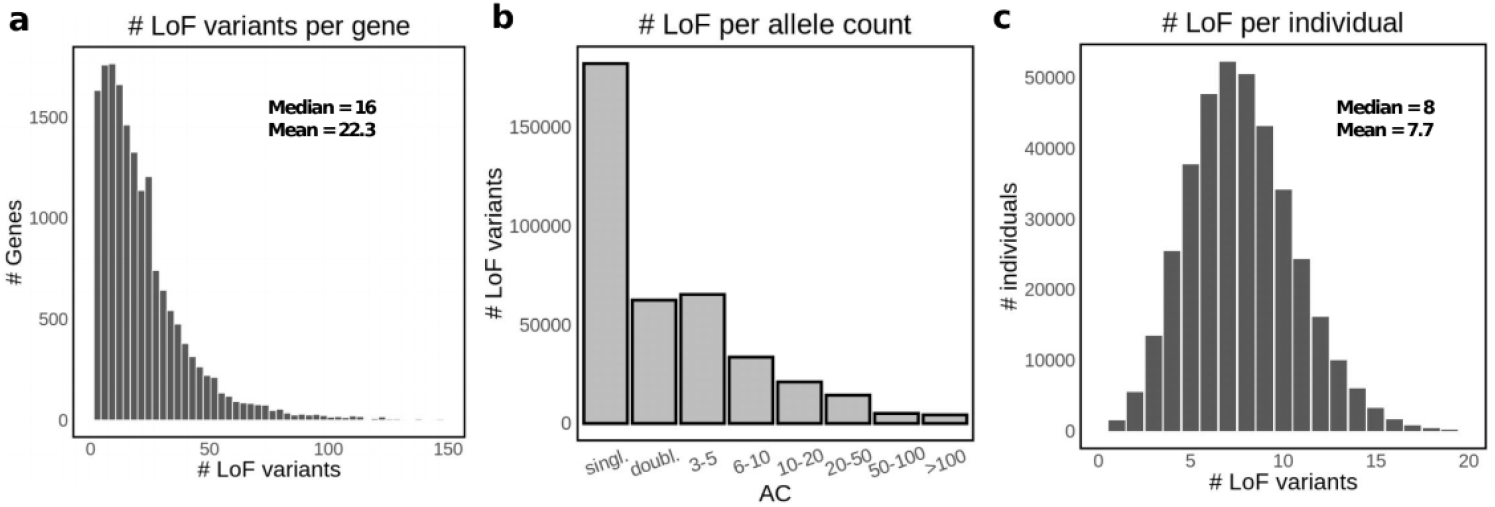
Frequency of loss-of-function (LoF) variants in the UK Biobank. (**a**) Number of LoF variants per protein-coding gene. 69 outlier values were excluded. (**b**) Number of LoF variants per allele count bin (N = 383,637). (**c**) Number of LoF variants per UK Biobank individual (N = 374,826). 11 outlier values were excluded.

**Supplementary Figure 11:**
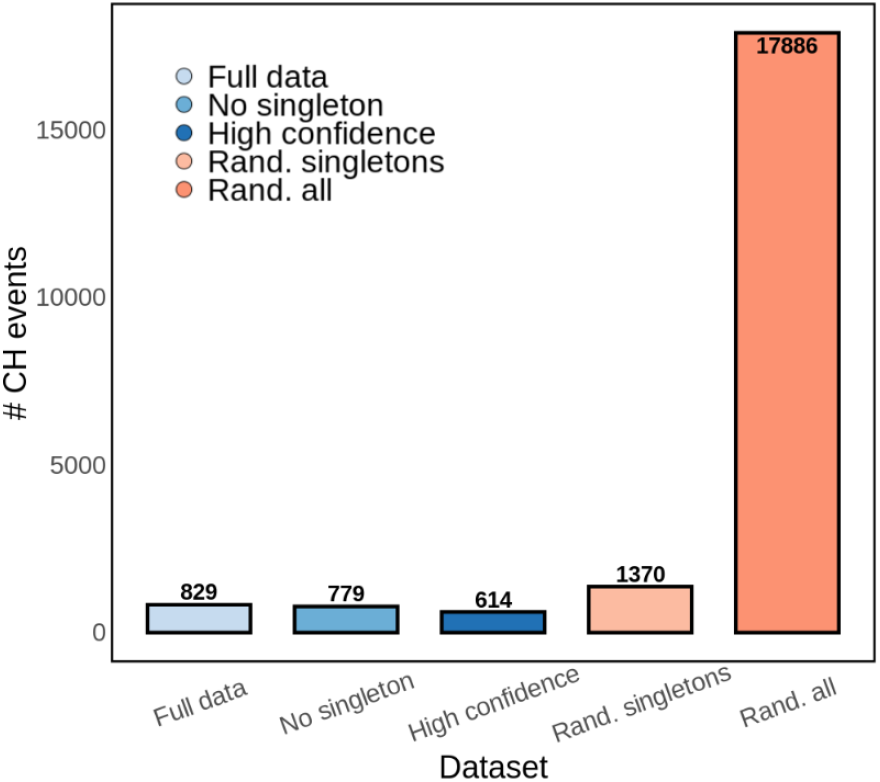
Number of compound heterozygous (CH) events across several categories. “Full data”: all LoF variants in the study, “No singleton”: all LoF variants except singletons, “High confidence”: LoF variants excluding singletons and calls with phasing confidence score < 0.99, “Rand. singletons”: including all LoF variants but shuffling phasing of singletons, “Rand. all”: shuffling phasing of all LoF variants.

**Supplementary Figure 12:**
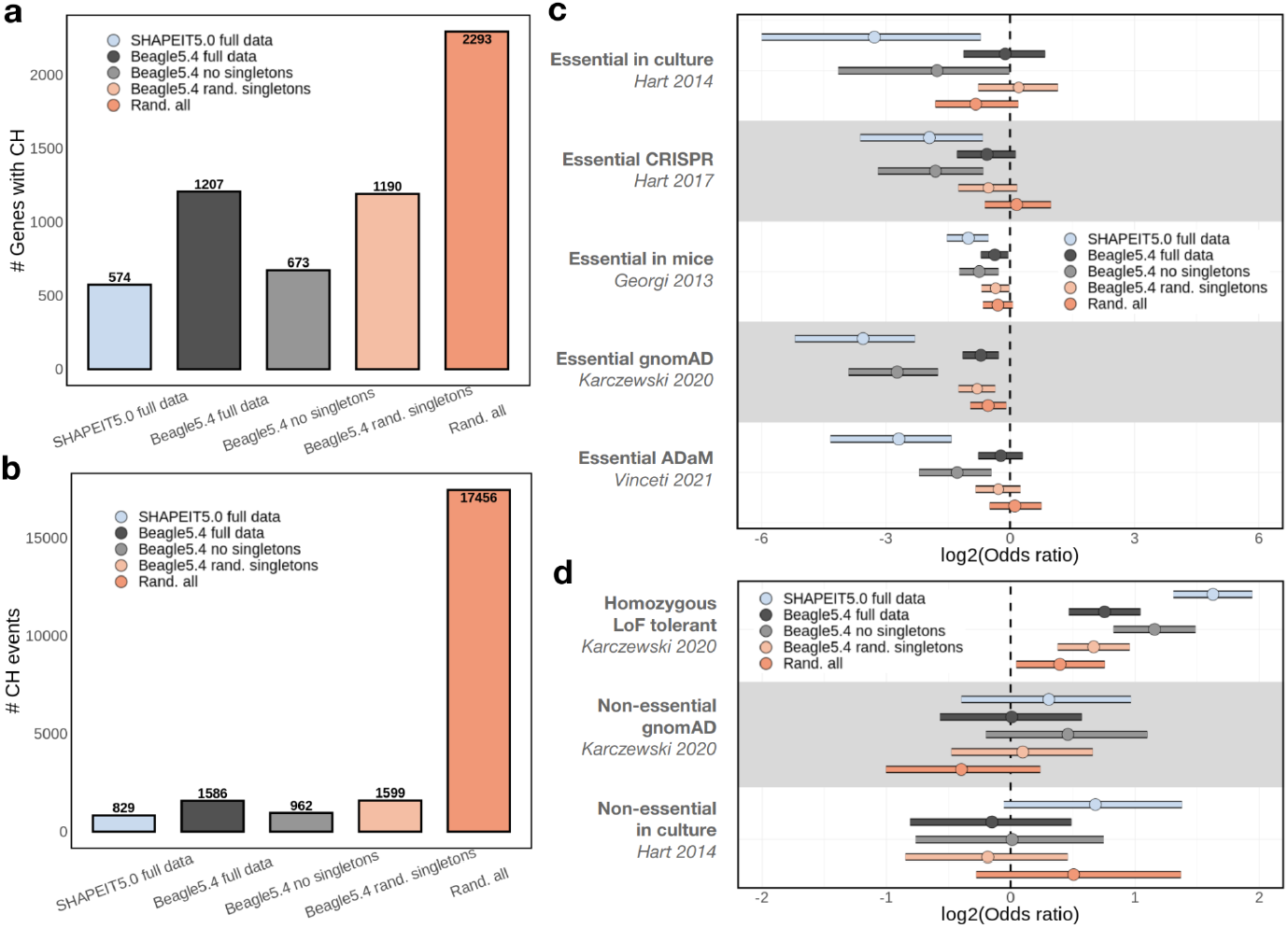
Compound heterozygous identification in the UK Biobank WES data phased with Beagle5.4. (**a**) Number of genes with a CH event in at least one individual. (**b**) Number of compound heterozygous (CH) events. (**c**) Two-way Fisher’s Exact test odds ratios and 95% confidence interval (log2-scaled) of CH genes versus non-CH genes presence in multiple lists of essential genes (see Methods). Background is composed of genes with ≥2 LoF mutations. X-axis is capped at -6. (**d**) Same as previous, but for across lists of non-essential genes.

**Supplementary Figure 13:**
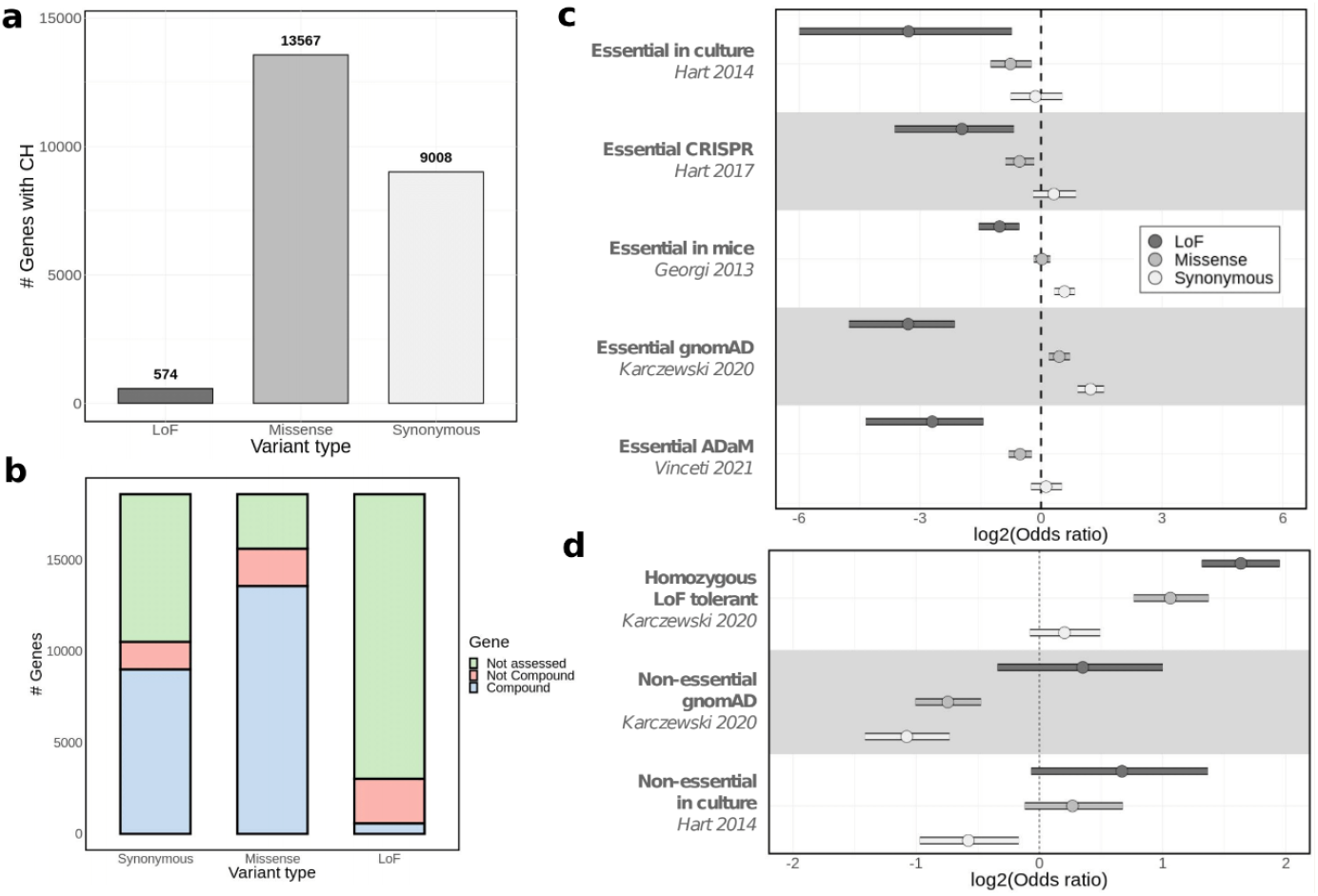
Comparison of compound heterozygous results with LoF, missense or synonymous variants. (**a**) Number of genes with a CH event in at least one individual. (**b**) From a total of 18,595 protein coding genes, number of genes with CH event (blue), number of genes with individuals with ≥2 mutations but found only in one haplotype (red), number of genes for which individuals with ≥2 mutations did not occur (green). (**c**) Two-way Fisher’s Exact test odds ratios and 95% confidence interval (log2-scaled) of CH genes versus non-CH genes presence in multiple lists of essential genes (see Methods). Background is composed of genes with ≥2 LoF mutations. X-axis is capped at -6. (**d**) Same as previous, but for across lists of non-essential genes.

**Supplementary Table 1.**
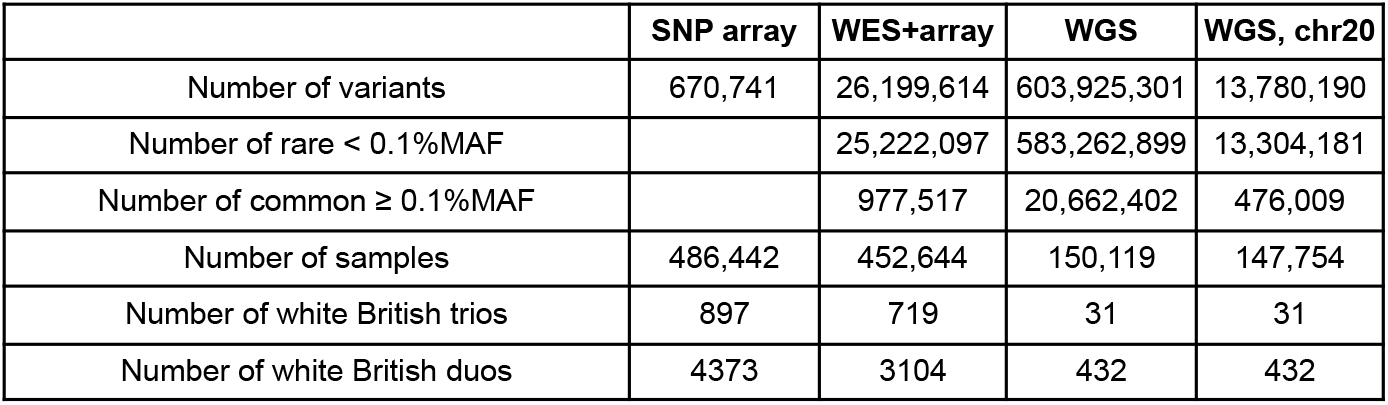
Summary statistics of the phased datasets.

|  | SNP array | WES+array | WGS | WGS, chr20 |
| --- | --- | --- | --- | --- |
| Number of variants | 670,741 | 26,199,614 | 603,925,301 | 13,780,190 |
| Number of rare < 0.1%MAF |  | 25,222,097 | 583,262,899 | 13,304,181 |
| Number of common ≥ 0.1%MAF | | 977,517 | 20,662,402 | 476,009 |
| Number of samples | 486,442 | 452,644 | 150,119 | 147,754 |
| Number of white British trios | 897 | 719 | 31 | 31 |
| Number of white British duos | 4373 | 3104 | 432 | 432 |

**Supplementary Table 2.** Running times and cost estimates for phasing WGS data with SHAPEIT5 or Beagle5.4 on the Research Analysis Platform (RAP) of the UK Biobank.

| Software | Step | Total time | Time per job | Virtual machine | Cost on-demand | Cost on-spot |
| --- | --- | --- | --- | --- | --- | --- |
| Beagle5.4 | Phasing | 24:45:00 | NA | mem3_ssd1_v2_x64 | 57.82 | 11.56 |
| Beagle5.4 | Indexing | 04:10:00 | NA | mem3_ssd1_v2_x8 | 0.96 | 0.19 |
| Beagle5.4 | Total chr20 | 28:55:00 | NA | NA | 56.86 | 11.37 |
| Beagle5.4 | Total whole genome | 60d, 5:50:00 | NA | NA | 2842.88 | 568.58 |
| SHAPEIT5 | Phasing common variants | 47:14:00 | 02:57:00 | mem3_ssd1_v2_x32 | 55.17 | 11.03 |
| SHAPEIT5 | Ligation | 01:24:00 | 01:24:00 | mem3_ssd1_v2_x2 | 0.10 | 0.02 |
| SHAPEIT5 | Phasing rare variants | 16:33:00 | 01:02:00 | mem3_ssd1_v2_x32 | 19.33 | 3.87 |
| SHAPEIT5 | Concatenating | 04:28:00 | 04:28:00 | mem3_ssd1_v2_x2 | 0.33 | 0.07 |
| SHAPEIT5 | Total chr20 | 69:39:00 | 09:51:00 | NA | 74.93 | 14.99 |
| SHAPEIT5 | Total whole genome | 145d, 2:30:00 | 20d, 12:30:00 | NA | 3746.36 | 749.272 |

## Notes

### Competing Interest Statement

The authors have declared no competing interest.

### Summary of Updates

Supplementary Figure 7 is revised.

